# Statistical end-to-end analysis of large-scale microbial growth data with DGrowthR

**DOI:** 10.1101/2025.03.25.645164

**Authors:** Medina Feldl, Roberto Olayo-Alarcon, Martin K. Amstalden, Annamaria Zannoni, Stefanie Peschel, Cynthia M. Sharma, Ana Rita Brochado, Christian L. Müller

**Affiliations:** Department of Statistics, Ludwig-Maximilians-Universität München, Munich, Germany; Institute of Computational Biology, Helmholtz Munich, Munich, Germany; Department of Microbiology, Biocenter, Julius-Maximilians-Universität Würzburg, Würzburg, Germany; Interfaculty Institute of Microbiology and Infection Medicine Tübingen (IMIT), University of Tübingen, Tübingen, Germany; Cluster of Excellence ‘Controlling Microbes to Fight Infections’ (CMFI), University of Tübingen, Tübingen, Germany; Department of Molecular Infection Biology II, Institute of Molecular Infection Biology (IMIB), Julius-Maximilians-Universität Würzburg, Würzburg, Germany; Center for Computational Mathematics, Flatiron Institute, New York, USA; Microbial Automation and Culturomics Core Facility, European Molecular Biology Laboratory, Heidelberg, Germany

## Abstract

Quantitative analysis of microbial growth curves is essential for understanding how bacterial populations respond to environmental cues. Traditional analysis approaches make parametric assumptions about the functional form of these curves, limiting their usefulness for studying conditions that distort standard growth curves. In addition, modern robotics platforms enable the high-throughput collection of large volumes of growth data, thus requiring strategies that can analyze large-scale growth data in a flexible and efficient manner. Here, we introduce `DGrowthR`, a statistical R framework and standalone app with a no-code interface for the integrative analysis of large growth experiments. `DGrowthR` comprises methods for data pre-processing and standardization, exploratory functional data analysis, and non-parametric modeling of growth curves using Gaussian Process regression. Importantly, `DGrowthR` includes a rigorous statistical testing framework for differential growth (DG) analysis. To illustrate the range of application scenarios of `DGrowthR`, we analyzed three large-scale bacterial growth datasets targeting distinct scientific inquiries. On an in-house dataset comprising more than 20, 000 growth curves of two pathogens that were subjected to chemical perturbations, `DGrowthR` enabled the discovery of compounds with *significant* growth inhibitory effects as well as compounds that induce non-canonical growth dynamics. On two publicly available perturbation datasets (> 100, 000 growth curves), DG analysis recovered reported adjuvants and antagonists of antibiotic activity, as well as bacterial genetic factors that determine susceptibility to specific antibiotic treatments. We anticipate `DGrowthR` to streamline the analysis of high-volume growth experiments, enabling researchers to make biological discoveries in a standardized and reproducible manner.

## Introduction

The quantitative analysis of microbial growth is essential for characterizing bacterial populations. Experiments where measurements of bacterial growth (typically reported as optical density (OD)) are taken at multiple time points provide valuable information about the dynamics of bacterial populations over time [1, 2]. Current automatic platforms enable researchers to collect large volumes of these growth curves with ease by gathering data from multiple samples simultaneously [3]. The readouts from these experiments have the potential to provide valuable insights into the mechanisms underlying the bacterial response to various environmental and chemical cues. Appropriate statistical approaches and computational tools are necessary in order to gain valuable insights from these high-throughput data.

Traditional approaches to model time-series growth data, such as the Logistic model, make parametric assumptions about the functional form of these curves [4]. Multiple popular software packages, including `Sicegar` [5] and `Growthcurver` [6], provide options for modeling single growth curves with sigmoidal or double sigmoidal fits and extracting relevant microbial growth parameters. However, growth curves gathered under various environmental challenges can deviate significantly from these canonical forms and therefore violate the parametric assumptions of these models. While more flexible approaches for growth curve modeling exist, including `grofit` [7], `pyphe` [8], `gcplyr` [9], and `QurvE` [10], they often rely on techniques such as splines and are sensitive to outliers and technical variations. Furthermore, the majority of tools lack principled statistical testing capabilities for comparing growth curve characteristics across different experimental conditions. Notably, `NeckaR` [3] can compare growth curves based on area under the curve (AUC) values, but does not extract relevant growth parameters. Finally, one of the more comprehensive solutions toward growth curve modeling is offered by `Kinbiont` [11], a Julia-based framework that includes a host of microbial kinetics models for data fitting, several data analysis capabilities, and interpretable machine learning tools for downstream analysis and hypothesis generation.

To model and analyze more complex bacterial growth dynamics, Gaussian Process (GP) regression has recently emerged as a valuable alternative [12–14]. GP regression is a non-parametric approach that does not make explicit assumptions about the functional form of the growth curves and can jointly model multiple (replicate) growth curves, thus reducing the influence of outliers. The outcome of GP regression can be used to determine robust growth parameters, such as growth rates, carrying capacity, and AUC values, as well as determine different phases of growth [12, 14]. Additionally, Tonner et al. [13] showed that GP regression can be leveraged to perform hypothesis testing, comparing complete growth dynamics instead of relying on singular growth parameters. The Python package `AMiGA` [14] provides a comprehensive framework for the analysis of microbial growth curves using GP regression, yet requires the use of command-line instructions. Furthermore, none of the tools provide functionality to perform exploratory data analysis in the form of functional data analysis [15], clustering, or popular dimensionality reduction techniques.

In this contribution, we present `DGrowthR`, an R package and standalone desktop application, that enables statistical end-to-end analysis of high-throughput microbial growth datasets. `DGrowthR` offers functionality for data pre-processing, normalization, and standardization, and supports a wide range of visualizations for large-scale datasets. This includes both plotting tools for individual curves and low-dimensional embeddings of large curve collections via Functional Principal Component Analysis (FPCA) and Uniform Manifold Approximation and Projection (UMAP) [16–18]. Notably, these embeddings can facilitate the identification of distinct growth patterns (clusters) within the data by employing recent functional data clustering techniques [19]. Furthermore, `DGrowthR` can flexibly model collections of growth curves using GP regression and automatically extract all relevant growth curve characteristics. `DGrowthR` also includes a computationally efficient permutation-based statistical testing framework for comparing growth curves across different experimental conditions via Gamma-approximated permutation p-values [20, 21], speeding up computation by several orders of magnitude. The modular software structure of `DGrowthR` also facilitates storing of all relevant analysis results in structured objects that can be conveniently accessed for custom downstream analysis.

We illustrate the broad applicability and the unique features of `DGrowthR` by analyzing three large-scale datasets that comprise different bacterial species and their response to various forms of genetic and chemical perturbations. The first dataset consists of in-house chemical screens of two pathogens, *Salmonella enterica* and *Campylobacter jejuni*, against a diverse chemical library of 2,415 compounds. The second dataset is from Brenzinger et al. [22], where changes in the susceptibility of *Vibrio cholerae* to various chemical challenges are measured upon deletion of the cyclic-oligonucleotide-based anti-phage signaling system (CBASS). The third dataset from Brochado et al. [23] comprises measurements to study the effect of pairwise drug combinations on the growth of gram-negative pathogens. Using `DGrowthR`, we not only recapitulate the results of the original studies in an automated and reproducible fashion but also uncover additional insights into the growth dynamics of the bacteria in the different scenarios. We posit that `DGrowthR` provides a general, flexible statistical end-to-end framework, enabling microbiologists to streamline their analysis of large-scale microbial growth datasets in a computationally efficient and reproducible fashion.

## Results

### The DGrowthR framework

`DGrowthR` is a comprehensive R package and standalone desktop application designed for the analysis of high-throughput growth experiments. As such, it offers functionality for reading multiple data files from automatic plate readers as well as any metadata associated, such as experimental conditions. A commonplace example of such high-volume experiments includes chemical library screenings (Figure 1a). The resulting data is stored in a custom object, which conveniently stores all subsequent analysis results.

**Figure 1:**
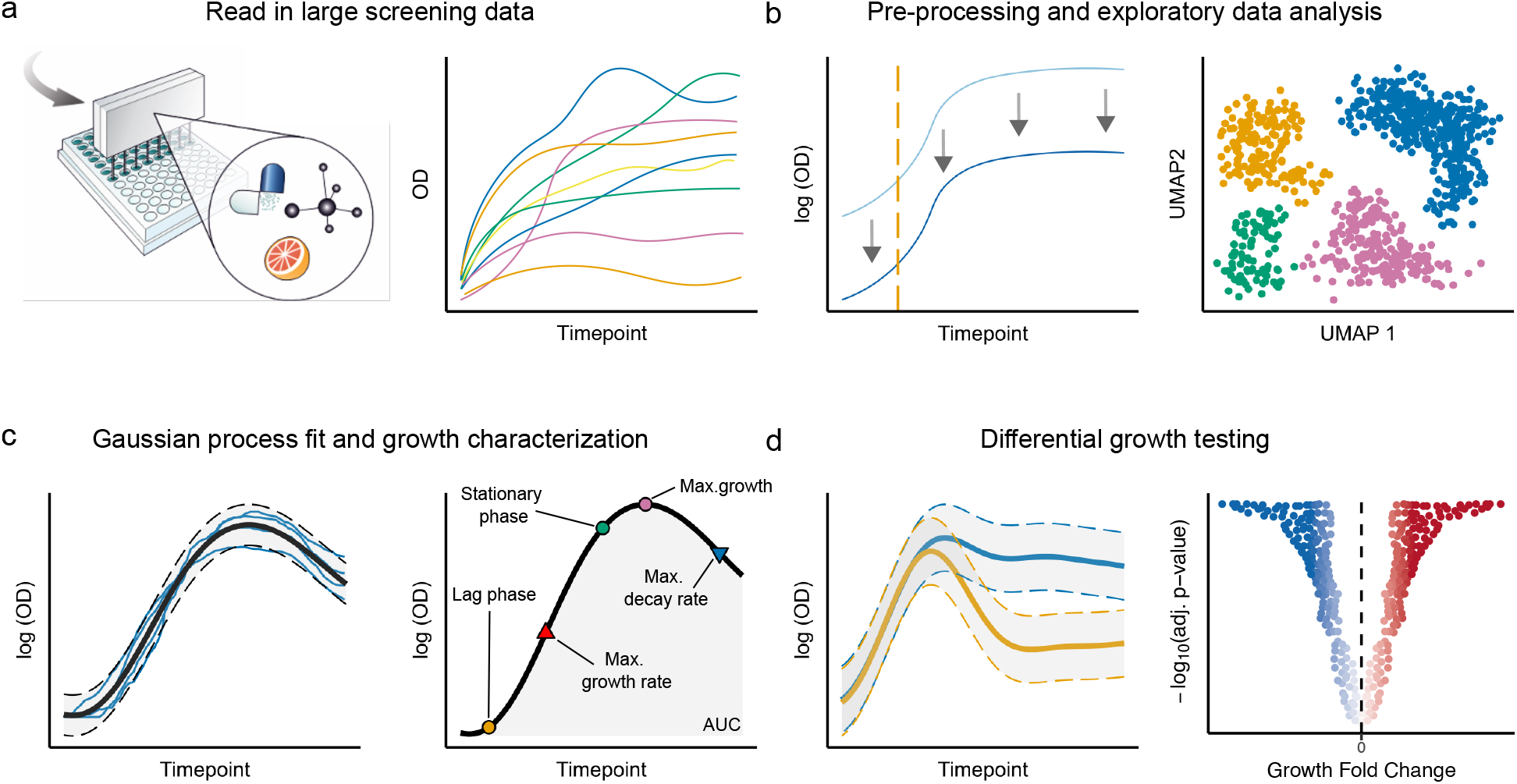
Schematic overview of the DGrowthR software framework. **a.** DGrowthR can read-in data from automatic plate-readers that gather data for multiple growth curves simultaneously. These curves can be visually inspected to decide on the next steps. **b**. Raw growth curve pre-processing includes baseline correction, log-transformation of OD measurements, and determination of measurement starting points. Exploratory data analysis via low dimensional embeddings includes FPCA and UMAP, followed by density-based clustering of the different growth dynamics. **c**. GP regression is used to flexibly model growth curves and extract relevant growth parameters, including maximum growth and decay rates and Area Under the Curve (AUC) values. Replicate growth curves can be pooled and modeled jointly. **d**. GP regression is combined with permutation-based testing to compare growth curves from different experimental conditions, providing statistical evidence for differential growth (DG).

The raw OD measurements can be visually inspected in order to determine appropriate pre-processing steps (Fig. 1a). The DGrowthR package includes common pre-processing functions including the removal of initial time points (typically due to noisy measurements), log-transformation of OD measurements, and baseline correction such that every growth curve’s initial measurements start at zero. The pre-processed growth curves can be visually inspected (Figure 1b). To characterize the different growth dynamics present in this large volume of data, `DGrowthR` offers functionality for embedding growth curves into a low-dimensional space using Functional Principal Component Analysis (FPCA) and Uniform Manifold Approximation and Projection (UMAP). Density-based clustering of these embeddings (such as with DBSCAN) clusters growth curves with similar dynamics and identifies potential outliers (Figure 1b) [16, 17]. The quantitative analysis of growth curves is performed using Gaussian Process (GP) regression, a non-parametric approach that can model complex growth dynamics without making assumptions about the functional form of the growth curves. GP regression is used to model each growth curve, individually or by pooling replicates. The resulting model can be directly used to quantify growth parameters such as maximum growth, area under the curve (AUC), and growth loss (Figure 1c). The first derivative of the model is leveraged to identify maximum growth and decay rates as well as doubling time. The second derivative is used to identify the moments of greatest increase and decrease in the growth rate which, in the case of sigmoidal-shaped growth curves, correspond to the end of the lag phase and the start of the stationary phase of growth, respectively (Supplementary Figure 2). These growth parameters can also be determined by sampling from the posterior of the resulting growth model. Finally, `DGrowthR` provides a statistical testing framework for comparing growth curves from different experimental conditions [13]. This framework is based on permutation testing, where the labels of the growth curves are permuted, and the Bayes Factor between the alternative and null models is calculated. To enable accurate and computationally efficient approximation of small p-values, we can limit the number of necessary permutations by approximating the distribution of the test statistics with a Gamma distribution [20, 21]. We adjust the resulting p-values for multiple comparisons using the Benjamini-Hochberg procedure [24], thus controlling the False Discovery Rate (FDR) to determine the statistical significance of the observed differences in growth dynamics (Fig. 1d). This provides an evidence-based approach to compare growth curves that takes the behavior of the entire growth dynamic into account, rather than individual growth parameters or other summary statistics. The overall R package workflow diagram for `DGrowthR` is available in Supplementary Fig. 1, detailing all available functions and corresponding input-output behavior. The `DGrowthR` repository is available at https://bio-datascience.github.io/DGrowthR/.

We next showcase the capabilities of `DGrowthR` by analyzing three independent datasets, dealing with different bacterial species and their response to various forms of chemical perturbations.

### Exploratory data analysis of large chemical screens with DGrowthR

In recent years, there has been a growing body of evidence showing that the growth of bacteria can be significantly altered by exposure to various types of chemical compounds, including non-antibiotic drugs [25–28]. The susceptibility of bacteria to these compounds varies depending, among many other factors, on the species in question [25, 28, 29]. Here, we analyze an in-house screen of two gram-negative pathogens, *Salmonella enterica* serotype Typhimurium (*S. enterica*) and *Campylobacter jejuni* (*C. jejuni*), against a diverse chemical library of 2,415 compounds from MedChemExpress. This library comprises FDA-approved drugs, metabolites, and food-additive homologous compounds. The growth of bacterial species was monitored using a high-throughput plate reader, capturing OD measurements at regular intervals (**Methods**). The resulting dataset contains 16,128 growth curves for *S. enterica* and 5,376 for *C. jejuni*, with 17 time points per growth curve in each instance.

To provide a global overview of these large-scale screens, `DGrowthR` encompasses low-dimensional embeddings and automatic clustering of growth curves (Fig. 2). Functional PCA (FPCA) (Fig. 2a and d) reveals that, for both organisms, *>* 99% of their growth curve variability can be captured in the first two functional principal components (FPC-1 and FPC-2) where growth curves (represented as points in the embedding) are forming a contiguous transition from non-increasing curves (colored in blue) to moderately increasing (colored in orange) and strongly increasing curves (green). `DGrowthR`’s UMAP representation, conversely, separates the non-increasing and moderately increasing curves from the more regular growth curves, allowing `DGrowthR`’s subsequent density-based clustering routine [19] to identify non-trivial clusters. Figure 2 c and f visualize the collection of growth curves according to their cluster membership. Cluster 1 (colored in green) represents increasing, cluster 2 (colored in blue) non-increasing, and cluster 0 (colored in orange) moderately increasing lagged growth curves.

**Figure 2:**
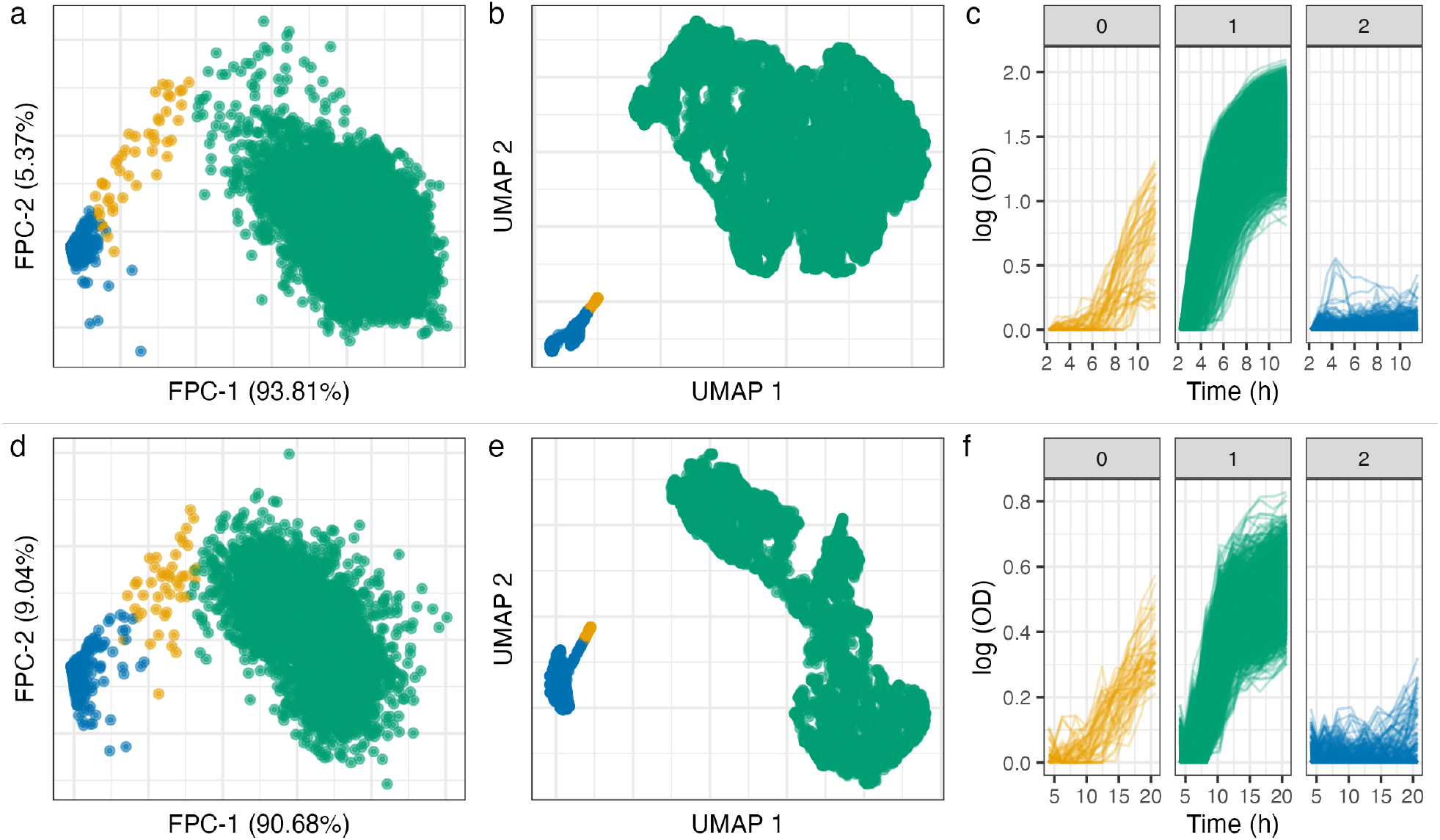
Exploratory analysis of a large chemical screen. **a.***S. enterica* growth curves (n=16,128) embedded in the first two FPCs. The percentage of the total variance explained by each FPC is shown in parenthesis. Points are colored based on cluster membership (see panel **c**). **b**. UMAP embedding of *S. enterica* growth curves, colored by cluster membership. UMAP coordinates are used to cluster growth curves with DBSCAN. **c**. Growth curves for *S. enterica* colored by cluster membership. **d**. *C. jejuni* growth curves embedded in the first two FPCs (see panel **f**) (n=5,376). **e**. UMAP embedding of *C. jejuni* growth curves (n=5,376), colored by cluster membership. UMAP coordinates are used to cluster growth curves with DBSCAN . **f**. Growth curves for *C. jejuni* colored by cluster membership.

Overall, this analysis indicates (i) that exposure to chemical stress gives rise to a variety of growth dynamics for *S. enterica* and *C. jejuni*, beyond a simple binary classification of growth or no growth, and (ii) that `DGrowthR` can visualize and categorize the different dynamics consistently across the two species.

### Flexible growth curve and parameter estimation with DGrowthR

We next applied `DGrowthR` to model the underlying growth response of *S. enterica* and *C. jejuni* to each chemical challenge. GP regression is leveraged to pool all replicate growth curves that correspond to the same treatment. After GP model fitting, we estimate all relevant growth parameters from the first and second derivatives of the GP (see **Methods** and Supplementary Fig. 2). To illustrate the necessity of GP-based model estimation for these data, we compared the quality of the fits with parametric growth curve estimation using `Growthcurver` [6] on the complete *S. enterica* dataset. We observed that (i) `Growthcurver` is unable to fit the majority of non-increasing curves (Supplementary Fig. 3a-c) and (ii) `Growthcurver` provides parametric curve fits to data with non-canonical growth dynamics (Supplementary Fig. 3d-f) that do not capture the true underlying dynamic, as compared to `DGrowthR` (Supplementary Fig. 3g-i). Blind large-scale application of such parametric modeling would thus result in inaccurate estimation of growth parameters, severely hampering downstream analysis.

We next illustrate `DGrowthR`’s capabilities to provide global visualizations of the derived growth parameters. Figure 3 shows the estimated changes in AUC and maximum growth rate with respect to the values obtained for control DMSO treatments for each compound and species, respectively. We observed that, in the case of *S. enterica*, the majority of treatments result in a decrease of AUC and maximum growth rate (*n* = 1191, 49.3%, Fig.3a, lower left quadrant), with a comparable fraction (*n* = 1155, 47.8%) showing higher AUC but lower maximum growth rate compared to the DMSO. Only 2.4% (*n* = 59) of compounds increased both parameters and 0.3% (*n* = 8) decreased AUC while increasing maximum growth rate. Coloring the estimated growth parameters according to (pooled) cluster membership of the underlying growth curves also confirms that they recapitulate the observed growth patterns in Fig.2. `DGrowthR` also allows annotation of individual treatment, as exemplified by the compound Paromomycin, an anti-parasitic drug, that induces a reduced maximum growth rate and AUC fold change (Fig. 3a). For the *C. jejuni* screens, we observed similar overall effects of the chemical challenges on the respective growth parameters, although with considerably higher diversity in the upper quadrants (Fig.3b). While the lower-left quadrant still contained the largest group (*n* = 1142, 47.3%), treatments increasing AUC with lower maximum growth rate accounted for 37.6% (*n* = 908), and notably more compounds increased maximum growth rate compared to *S. enterica* (*n* = 171, 7.1% and *n* = 159, 6.6% for higher and lower AUC, respectively). For instance, the antipsychotic drug Droperidol results in a 1.26-fold increase in AUC yet a lower maximum growth rate with respect to DMSO.

**Figure 3:**
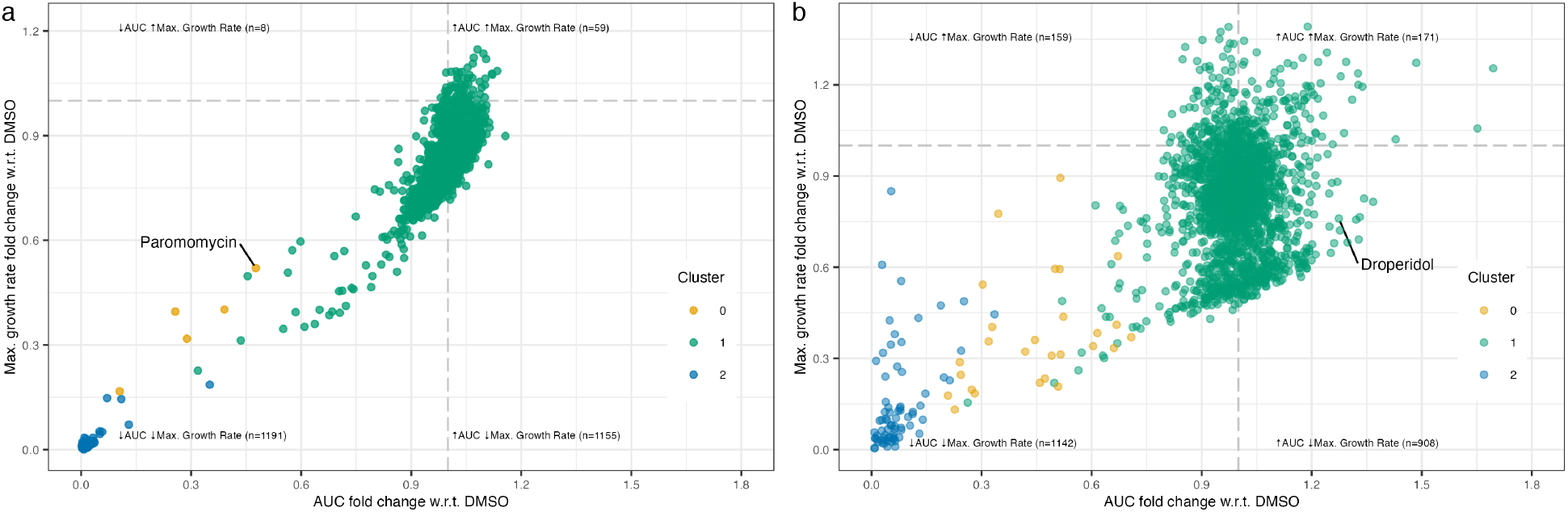
Visualization of estimated growth parameters with DGrowthR. All replicate growth curves are pooled and modeled with GP regression. Growth parameters are determined from the resulting model. **a.**Fold changes in AUC and maximum growth rate for each compound relative to DMSO control wells, shown for *S. enterica*. **b**. Fold changes in AUC and maximum growth rate for each compound relative to DMSO control wells, shown for *C. jejuni*. In both panels, points are colored according to cluster membership from Fig.2.

In summary, `DGrowthR`’s ability to flexibly model the whole range of observed growth dynamics facilitates a comprehensive and consistent estimation of derived growth parameters and enables direct comparison between different large-scale screens.

### Differential growth (DG) analysis with DGrowthR

While investigating individual growth characteristics and parameters is informative, summarizing and comparing growth curves across multiple growth metrics can be cumbersome for large-scale screens. Moreover, since the estimation of growth parameters can be sensitive to the experimental setup, the magnitude and scale of the modeled growth curves can differ substantially across experiments [13, 14].

To alleviate these shortcomings, `DGrowthR` includes a rigorous statistical testing framework that enables differential growth (DG) estimation across the *entire* growth curve. Briefly, we compare growth curves from exposure to a given compound with those from control conditions. For the present screens, we use DMSO treatment as the control condition. Growth curves are modeled using two GP regression models: the null model, in which all growth curves are pooled and modeled only as a function of time, and the alternative model, where growth curves are modeled as a function of time and treatment. The marginal likelihoods of the data under each model are calculated, and the ratio of these values represents the observed Bayes Factor. To determine the statistical significance of the observed Bayes Factor, we permute the treatment labels of the growth curves to obtain a null distribution of the Bayes factor test statistic. A Gamma-approximated p-value (p-value_gam_) is then calculated as the proportion of permutations that result in a Bayes factor greater than the observed value (see **Methods**). In this way, we determine the statistical significance of the observed differences in growth dynamics by examining complete growth curves, rather than comparing individual growth parameters.

Figure 4a and b illustrate the results of `DGrowthR`’s DG analysis for *S. enterica* and *C. jejuni*, respectively. At a significance threshold of adjusted p-value_gam_ *<* 0.01, 115 compounds showed a significant effect on the growth of *S. enterica*, all exhibiting decreased growth with respect to DMSO (Figure 4a). As an illustrative example, Figure 4c shows the growth curves and associated GP model for the antirheumatic drug Auranofin (colored in blue) compared to DMSO (colored in grey). While Auranofin did not completely inhibit the growth of *S. enterica*, the resulting growth dynamic was deemed significantly different from the control by `DGrowthR`’s DG procedure. To highlight the statistical validity of `DGrowthR`’s testing framework, we show the Gamma-approximated p-value distribution for the *S. enterica* screen in Figure 4e. As expected, the p-value distribution is nearly uniform across the [0, 1] interval, with an enrichment for small p-values.

**Figure 4:**
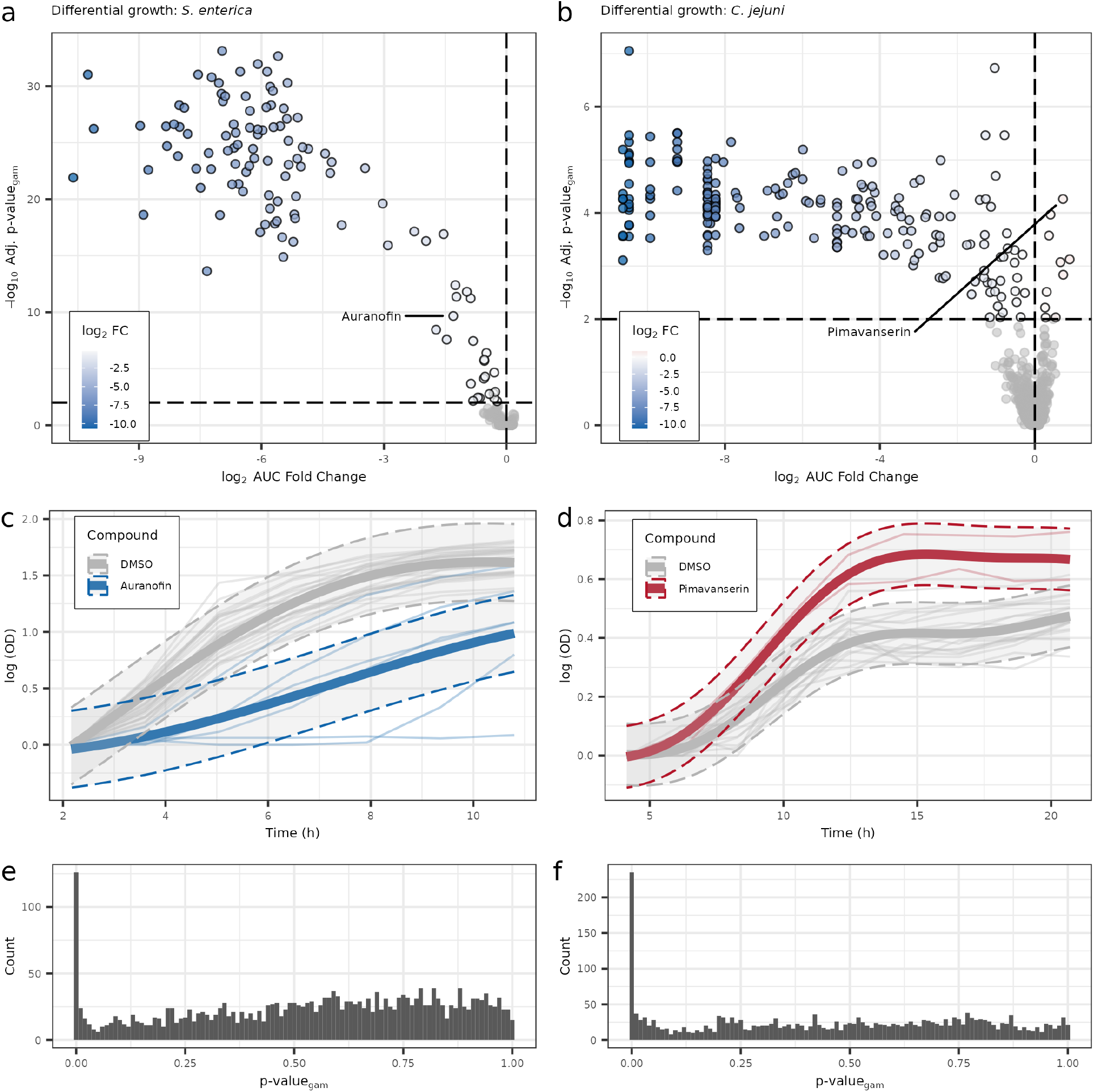
Differential growth analysis with `DGrowthR`. **a.**Volcano plot showing the results of the differential growth analysis for *S. enterica* against 2,415 compounds. Corrected gamma-approximated p-values (p-value_gam_) are compared to log_2_ fold change in AUC. The dashed horizontal line indicates an adjusted p-value_gam_ threshold of 0.01, while the vertical line indicates an absolute AUC log_2_ fold change of 0. **b**. Volcano plot showing the results of the differential growth analysis for *C. jejuni* against 2,415 compounds. Vertical and horizontal lines indicate the same values as in panel **a. c**. Alternative model of *S. enterica* growth treated with Auranofin (log_2_FC = −1.3, adjusted p-value_gam_ = 1.018 × 10^−9^). The shaded area indicates 95% confidence interval of the fitted model. Replicate growth curves under Auranofin treatment (n=6) and DMSO treatment (n=24) are shown. **d**. Alternative model of *C. jejuni* growth treated with Pimavanserin (log_2_FC = 0.73, adjusted p-value_gam_ = 0.000117). Replicate growth curves under Pimavanserin treatment (n=2) and DMSO treatment (n=18) are shown. **e**. Histograms of unadjusted gamma-approximated p-values for *S. enterica*. **f**. Histograms of unadjusted gamma-approximated p-values for *C. jejuni*.

In the case of *C. jejuni*, while the majority of treatments resulted in decreased growth (207 compounds), `DGrowthR` identified ten drugs that significantly increased the overall growth of *C. jejuni* (Figure 4b). One such compound was Pimavanserin, an antipsychotic drug. Indeed, Figure 4d depicts the growth curves gathered for *C. jejuni* treated with Pimavanserin (colored in red), which exhibited a higher AUC with respect to the control. Finally, the Gamma-approximated p-value distribution for the *C. jejuni* screen is shown in Figure 4f, confirming the validity of the testing procedure for this screen.

Together, these findings add to the growing body of evidence that specific drugs can significantly affect the growth dynamics of pathogenic bacteria. In our in-house large-scale screens, we found *C. jejuni* to be affected by a greater number of these compounds. We observed that the effects on growth can have a complex pattern, beyond simple growth inhibition, but can be reliably detected using `DGrowthR`. Further studies are required to investigate the mechanisms underlying these growth phenotypes and to determine their clinical relevance.

### Exploring the effects of genetic factors on bacterial growth with DGrowthR

The species-dependent variability of growth dynamics in our in-house chemical screens suggests multiple potential mechanisms of action for the compounds in question that also depend on the genetic makeup of the respective bacterium. A prime example of such a genetic dependency has been recently found by Brenzinger et al. [22] where the activity of antifolate antibiotics has been shown to be influenced by the presence of the cyclic-oligonucleotide-based anti-phage signaling system (CBASS) in *Vibrio cholerae*. We used `DGrowthR` to re-analyze the underlying dataset, which comprises growth curves for wild-type and a CBASS-operon deleted (ΔCBASS) strain of *V. cholerae*. The two strains were screened against a diverse chemical library of 94 compounds at multiple concentrations, resulting in 2,304 growth curves.

`DGrowthR’s` exploratory data analysis of the growth curves revealed a larger variability of growth dynamics in this screen compared to the previous in-house screen (Supplementary Figure 4) with four identified clusters (see Figure 5a). We pooled all replicate growth curves gathered for the same genotype-compound-concentration combination and obtained growth parameters from the resulting GP models. We again verified that the GP regression framework enables an overall improvement in fitting quality over parametric models, including logistic and Gompertz models. Indeed, for the vast majority of treatments, GP regression obtains the lowest mean-squared error fit compared to parametric models (Supplementary Material E and Supplementary Figure 5).

**Figure 5:**
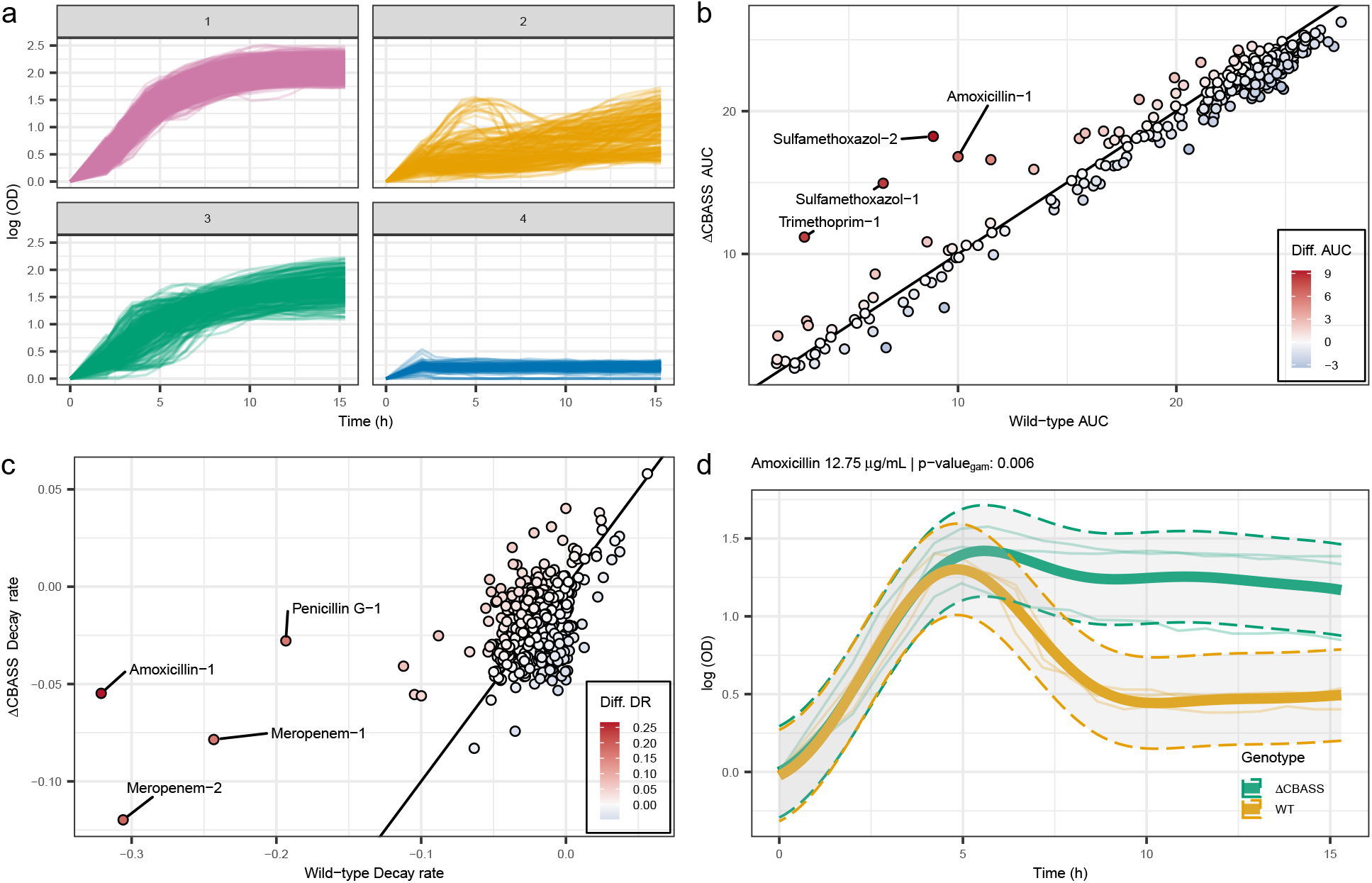
Analyzing the influence of the CBASS anti-phage defense system on antibiotic sensitivity with `DGrowthR`. **a.**All 2,304 growth curves from the chemical screen done by Brenzinger et al. [22] were embedded using FPCA and clustered into four groups based on DBSCAN. **b**. Comparison of AUC values determined by `DGrowthR`, for the wild-type and ΔCBASS strains of *V. cholerae* treated with the same compound-concentration combination. **c**. Comparison of maximum decay rates determined by `DGrowthR`, for the wild-type and ΔCBASS strains of *V. cholerae*. **d**. Alternative model of *V. cholerae* growth treated with Amoxicillin, comparing the wild-type and ΔCBASS strains. Shaded area indicates 95% confidence interval.

Mirroring Brenzinger et al.’s analysis, we used the AUCs of the estimated GP models to compare the effects of compounds on the wild-type strain and the ΔCBASS strain. Figure 5b shows the resulting AUC scatter plot, revealing the ΔCBASS strain exhibits increased AUC in the presence of Sulfamethoxazole and Trimethoprim, compared to the wild-type strain. This recapitulates the main findings by Brenzinger et al. where the ΔCBASS strain showed decreased susceptibility to Sulfamethoxazole and Trimethoprim, two antifolate antibiotics. Notably, `DGrowthR` also revealed that the ΔCBASS strain was less susceptible to Amoxicillin, an antibiotic that targets cell wall synthesis.

To showcase `DGrowthR`’s ability to analyze non-standard growth characteristics, we next investigated the differences between the wild-type and ΔCBASS strains in terms of *maximum decay rates* in response to the same chemical challenge. Figure 5c depicts a scatter plot of `DGrowthR`’s estimated maximum decay rates for the wild-type and ΔCBASS strains, revealing that the ΔCBASS strain showed a lower maximum decay rate in response to Amoxicillin, as well as Meropenem and Penicillin G, all known inhibitors of cell wall synthesis. Figure 5d shows the fitted growth curves for the wild-type and ΔCBASS strains treated with Amoxicillin (12.75 *µg*/mL), confirming the marked change in maximum decay rate.

In summary, our re-analysis of this dataset with `DGrowthR` adds further evidence to the noted influence of the CBASS system on the susceptibility to cell-wall synthesis inhibitor antibiotics, thus providing additional testable hypotheses for experimental determination of the underlying mechanisms.

### Exploring combinatorial treatment effects on bacterial growth with DGrowthR

Another common application of chemical screenings is to determine the effects of drug combinations on bacterial growth [23, 30]. For instance, Brochado et al. [23] measured the effects of almost 3,000 dose-resolved combinations of antibiotics, human-targeted drugs, and food additives on the growth of multiple gram-negative pathogens.

Here, we used `DGrowthR` to re-analyze a subset of these data, focusing on *Escherichia coli* BW (*E. coli*). In the original study, each drug combination was assessed in a 4 ×4 tailored-dose matrix (three concentrations per drug plus a no-drug control). Interactions were quantified using the Bliss independence model [31], which compares observed “fitness” values (relative growth compared to a reference) at a single time point (typically at the transition to stationary phase) between combinatorial treatment and the product of single treatments. With `DGrowthR`, we can complement this type of analysis by quantitatively assessing the significance of the effect of a combinatorial treatment compared to each individual treatment using DG testing along the entire growth curves. This can directly highlight particularly potent combinatorial effects, thus helping to prioritize interactions derived from Bliss models.

For systematic comparison, we focused on 78 unique compounds at their highest available concentrations, yielding a total of 13,610 growth curves. Due to the asymmetric plate design of the original screen, not all pairwise combinations were available. From the available data, we constructed 5,795 pairwise comparisons of *drug combinations* versus *individual treatments*, resulting in 2,867 unique two-compound pairs, each tested in both orientations (treatment A alone vs. A+B, and treatment B alone vs. A+B), producing 5,734 comparisons, with an additional 61 self-pair comparisons serving as controls.

To get a high-level summary of the complete Brochado et al. dataset, we visualized the FPCA embedding of all 115,668 growth curves in Fig. 6a (colored in grey) and the 13,610 *E. coli* BW growth curves colored by their maximum OD. This representation reveals a strong stratification by maximum OD along FPC-1 (94.5% explained variance), indicating that points with large separation along this principal component exhibit considerably different growth behaviors. As shown below, placing individual treatment curves alongside their corresponding combination curves in this space allows a direct assessment of potential synergistic or antagonistic effects. Large separations suggest substantial changes in growth dynamics upon combining drugs.

**Figure 6:**
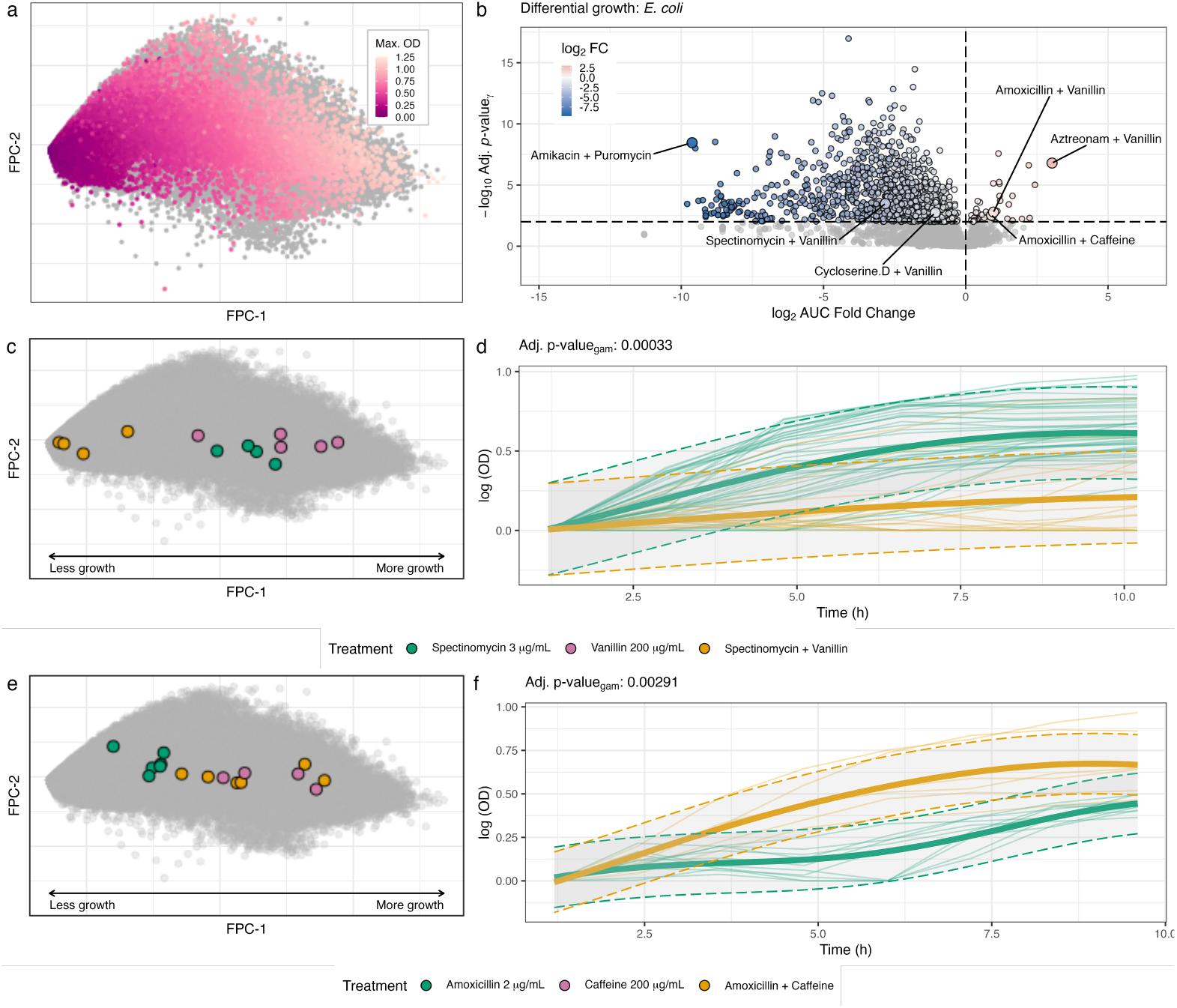
Analyzing the effects of pairwise drug combinations on the growth of *E. coli* BW with `DGrowthR`. **a.**FPCA embedding of all 115,668 growth curves from the chemical screen done by Brochado et al. [23] (colored in grey) and 13,610 *E. coli* BW curves (colored by maximum measured OD value) underlying the 5,795 tested drug combinations used in DG analysis. **b**. Volcano plot showing the results of the DG analysis of 5,795 pairwise drug interactions in *E*.*coli* BW using 500 permutations. Corrected gamma-approximated p-values (adjusted p-value_gam_) are compared to log_2_ fold change in AUC. The dashed horizontal line indicates an adjusted p-value_gam_ threshold of 0.01, while the vertical line indicates an absolute AUC log_2_ fold change of 0. Key validated interactions, including the Spectinomycin and Vanillin results and the Amoxicillin and Caffeine interaction, are explicitly labeled. **c**. Position of replicate growth curves in the FPCA embedding, gathered from treatment with Vanillin (200 *µg*/mL) and Spectinomycin (3 *µg*/mL) individually and in combination. **d**. Alternative model of *E. coli* BW growth treated with Spectinomycin alone and in combination with Vanillin. The shaded area indicates 95% confidence interval. Adjusted gamma approximated p-values after 500 permutations are shown. **e**. Position of replicate growth curves in the FPCA embedding, gathered from treatment with Amoxicillin (2 *µg*/mL) and Caffeine (200 *µg*/mL) individually and in combination. **f**. Alternative model of *E. coli* BW growth treated with Amoxicillin alone and in combination with Caffeine. The shaded area indicates 95% confidence interval. Adjusted gamma approximated p-values after 500 permutations are shown.

We next summarized our DG results on the *E. coli* BW subset in Fig. 6b, showing adjusted gamma-approximated p-values vs. log AUC fold changes. Out of the 5,795 tested combinations evaluated using 500 permutations and gamma-approximated p-values, 1,192 showed statistical significance (adjusted p-value_gam_ *<* 0.01). Among these, we successfully recovered key results from Brochado et al. [23], including the synergistic effect between Vanillin and the antibiotic Spectinomycin (DG for Spectinomycin + Vanillin in reference to Spectinomycin annotated in Fig. 6b). Figure 6c depicts the three corresponding sets of replicated growth curves, namely Spectinomycin (3 *µg*/mL) (green), Vanillin (200 *µg*/mL) (red), and the combination treatment (orange). Indeed, the location of the combination curves at small FPC-1 coordinates indicates growth inhibition, whereas the individual curves indicate normal growth. Figure 6d shows the corresponding GP fits of Spectinomycin and combined Spectinomycin-Vanillin treatment curves, respectively, and DG results (adjusted p-value_gam_ = 3.3 × 10^−4^).

Similarly, we identified a potential *antagonistic* effect of Caffeine (200 *µg*/mL) on treatment with Amoxicillin (2 *µg*/mL) (DG for Amoxicillin + Caffeine in reference to Amoxicillin annotated in Fig. 6b). The combination of Caffeine and Amoxicillin resulted in a higher maximum growth compared to Amoxicillin alone, though not as high as Caffeine alone (Fig. 6e). Figure 6f shows the corresponding GP fits of Amoxicillin and combined Amoxicillin-Caffeine treatment curves, respectively, and DG results (adjusted p-value_gam_ = 2.91 ×10^−3^), confirming the antagonistic effect. Interestingly, the effect of Caffeine against antibiotic treatment in *E. coli* has been validated and further investigated in an independent follow-up study [32].

More broadly, of the 1,192 statistically significant combinatorial effects, the vast majority (*n* = 1, 148) led to a negative AUC fold change, indicating that drug combinations typically resulted in lower growth than single treatment. The highest AUC fold change was observed for Aztreonam (0.06 *µg*/mL) and Vanillin (200 *µg*/mL) against the reference of only Aztreonam (log_2_FC = 3.03, adjusted p-value_gam_ = 1.59 ×10^−7^) (Supplementary Fig. 6a-b). Brochado et al. [23] also reported an antagonistic relationship for this pair, noting that Vanillin generally antagonizes many antibiotics. The lowest fold change was observed for the combination of Amikacin (0.75 *µg*/mL) and Puromycin (25 *µg*/mL) against Puromycin alone (log_2_FC = −9.62, adjusted p-value_gam_ = 3.47 ×10^−9^) (Supplementary Fig. 6c-d). This aligns with the findings in Brochado et al. [23], where synergies are significantly enriched within drugs with similar mechanisms of action. Additionally, the combination of D-Cycloserine (8 *µg*/mL) and Vanillin (200 *µg*/mL) showed a modest but significant reduction in growth compared to D-Cycloserine alone (log_2_FC = −1.07, adjusted p-value_gam_ = 1.78 ×10^−3^), further supporting the complex modulatory role of Vanillin in drug interactions (Supplementary Fig. 6e-f).

## Discussion

Here, we have introduced `DGrowthR`, an end-to-end framework for visualizing, modeling, and analyzing large-scale bacterial growth curve data. `DGrowthR` combines state-of-the-art statistical methods, including functional data analysis, Gaussian Process (GP) regression, and computationally efficient permutation-based inference schemes, into a comprehensive data analysis framework, enabling robust and reproducible growth data analysis. The successful use of GP regression in `DGrowthR` is consistent with previous work in microbial growth analysis, where GP models have been shown to offer flexibility beyond traditional parametric approaches [12–14]. Being available as fully documented R package and standalone desktop application, `DGrowthR` fills a critical software gap for microbial growth analysis since, e.g., existing R packages such as `gcplyr` and `NeckaR` are limited in their statistical tests, and packages such as `grofit` can struggle with non-sigmoidal growth patterns.

With `DGrowthR` we also put forward the idea that large-scale growth data from automated culturomics and screening platforms [3, 33] are amenable to statistical analysis in a similar fashion to data from, e.g., next-generation sequencing. Software packages (implementing dedicated statistical methods) such as `edgeR` [34] or `DESeq2` [35] have greatly advanced the field of transcriptomics by providing transparent and reproducible workflows for differential expression (DE) analysis. Using hundreds of thousands of bacterial growth curves from both in-house chemical screens and publicly available datasets [22, 23], we illustrated differential growth (DG) analysis workflows that can serve as templates for future end-to-end analysis of bacterial growth data.

In our in-house chemical screen, `DGrowthR` identified complex growth dynamics in *S. enterica* and *C. jejuni* exposed to different compounds. These findings are consistent with previous studies that have shown the potential of compounds to alter bacterial growth [25–28], and add new information for the studied pathogens. Among the compounds identified by DG analysis, Auranofin was found to significantly reduce the growth of *S. enterica*. This is in line with previous studies that have shown the potential of Auranofin as an antimicrobial agent, with reduced potency against Gram-negative species [36, 37]. In contrast, Pimavanserin was found to increase the growth of *C. jejuni*, which contrasts with the reported antibacterial activities of other antipsychotic drugs [26]. The re-analysis of the dataset from Brenzinger et al. [22] with `DGrowthR` confirmed the influence of the CBASS anti-phage defense system on the susceptibility of *V. cholerae* to antifolate antibiotics. With `DGrowthR`, we identified a decreased decay rate in response to cell wall synthesis inhibitors in the ΔCBASS strain, a relationship that can be further investigated to determine the underlying mechanisms.

`DGrowthR` analysis also reproduced one of the main findings from Brochado et al. [23] by confirming the synergistic effect between Vanillin and Spectinomycin, resulting in increased inhibition of *E. coli* growth. Additionally, we identified the antagonistic effect of Caffeine against the activity of Amoxicillin, leading to increased growth of *E. coli*. These findings were validated in an independent follow-up study [32]. DG analysis also enabled to highlight interactions that, to the best of our knowledge, have not been investigated beyond the original screen. DG analysis confirmed Vanillin as a broad antagonist for most antibiotic classes, an observation reported by Brochado et al. [23] but not followed up in that study or subsequent literature, with the Aztreonam-Vanillin combination exhibiting the highest positive AUC fold change among all tested pairs. The strongest synergistic effect was observed for the combination of Amikacin and Puromycin which has not yet been studied in the literature. Most notably, `DGrowthR` identified a novel interaction where combining Vanillin and D-Cycloserine resulted in a significant growth decrease, contrasting with the typically antagonistic behavior of Vanillin, thus suggesting a context-dependent modulatory role. These results demonstrate the ability of `DGrowthR` to detect understudied trends and new chemical interactions while providing a robust platform for generating testable hypotheses. Our results underscore the utility of `DGrowthR` in analyzing large-scale growth experiments in a diverse set of scenarios, where different research questions are prioritized. In all cases, the `DGrowthR` framework was able to provide novel insights into the growth dynamics of the studied bacteria and to suggest testable hypotheses for further investigation.

While `DGrowthR` addresses several existing limitations in microbial growth modeling and DG analysis, a number of challenges remain. Large data sets require extensive permutation testing, which can be computationally intensive and may limit accessibility to users without high-performance computing resources. Our approach of including Gamma approximations of p-values is a first step toward reducing the computational burden for the user. Nevertheless, the robustness of permutation tests could be compromised if there is a significant imbalance or only a small set of replicates available in the condition and control groups. This implies that `DGrowthR`’s hypothesis testing framework, like all permutation-based approaches, requires sufficiently many replicates to be powerful. While `DGrowthR` can model any curve shape, it currently only supports the identification of standard growth phases (lag, exponential, stationary). Ongoing work is concerned with the detection of repeated phase transitions, such as those arising from diauxic shifts [11, 14].

In summary, we believe that `DGrowthR` provides a valuable statistical framework for the analysis of largescale growth curve data. With the rapidly growing availability of robotics platform-based chemical screen protocols [3], we anticipate `DGrowthR` to play a substantial role in the robust and reproducible analysis of future large-scale microbial growth curve data.

## Methods

### Overview of the DGrowthR Framework

`DGrowthR` is an R package designed for the comprehensive analysis of bacterial growth curves. It efficiently handles high-dimensional data, integrating both growth measurements and metadata. By leveraging advanced statistical techniques, the package enables modeling and interpretation of growth dynamics under diverse experimental conditions.

### Data Input and Pre-processing

#### Data Structuring

`DGrowthR` supports direct import of data typically output by automated plate readers. The package accommodates multiple input files, aligning them along a common time axis for consistency. All imported data are stored in a structured object (a R S4 object) that also retains associated experimental metadata. To streamline analysis, `DGrowthR` utilizes an object-oriented approach, where a specialized object is created to hold both growth curve data and metadata. This structure consists of multiple slots that store intermediate analysis results, minimizing redundant computations and optimizing efficiency.

#### Pre-processing

Pre-processing is crucial for ensuring data quality and accurate modeling, particularly in large datasets derived from multiple experiments. `DGrowthR` provides several pre-processing options after data import, including the removal of initial time points, logarithmic transformation, and baseline correction. Early time points in a growth curve often contain technical noise and may not be informative; therefore, `DGrowthR` allows users to specify the number of initial time points to remove. Logarithmic transformation is applied to optical density (OD) measurements to better model exponential growth, facilitating the estimation of maximum growth rates and the identification of transition phases. Baseline correction addresses instrument fluctuations and non-biological variations in initial OD readings. Since the initial measured population size can vary across experiments, baseline correction ensures consistency by subtracting the initial population size from all subsequent measurements.

### Low-Dimensional Embedding and Visualization

`DGrowthR` is designed for detailed analysis of large-scale data sets. These often contain subtle growth patterns that may not be immediately apparent. To reveal these patterns, we implemented two low-dimensional embedding techniques: Functional Principal Component Analysis (FPCA) and Uniform Manifold Approximation and Projection (UMAP).

#### Functional Principal Component Analysis

Like Principal Component Analysis (PCA), Functional Principal Component Analysis (FPCA) aims to capture the primary sources of variation within the dataset for functional (e.g., time series) data. This approach allows users to identify dominant growth patterns within their data [38, 39]. In `DGrowthR`, FPCA calculations for growth curves are implemented using the `fdapace` package [40].

#### Uniform Manifold Approximation and Projection

Uniform Manifold Approximation and Projection (UMAP) provides non-linear dimensionality reduction, attempting to capture the intrinsic structure of complex datasets while projecting the data to a chosen low target dimension (typically two or three dimensions). This method facilitates visualization and highlights patterns or discrepancies in growth curves that might otherwise remain undetected in high-dimensional data [18]. Since individual UMAP coordinates do not possess an explicit meaning due to rotational invariance, `DGrowthR` aligns UMAP coordinates to match FPCA coordinates using the Kabsch algorithm [41]. After mean centering, the Kabsch algorithm calculates the optimal rotation matrix of points in UMAP coordinates that minimizes the root mean squared deviation to the points in FPCA coordinates. This makes UMAP visualization more interpretable.

#### Clustering

To further explore dataset structure, we implemented Density-Based Spatial Clustering of Applications with Noise (`DBSCAN`) on the UMAP embeddings [17]. `DBSCAN` identifies clusters based on local density and is particularly effective in handling noise and outliers [42, 43]. An adaptation of `DBSCAN` from Herrmann et al. [17] enhances cluster analysis by exploiting the topological structure found by UMAP. This strategy is implemented in `DGrowthR` using the `dbscan` package [44]. In the context of growth curves, this clustering can be used to identify groups of curves that exhibit similar growth dynamics.

FPCA and UMAP visualizations can be annotated with cluster assignments or metadata categories, enhancing the interpretability of growth patterns under different conditions.

### Gaussian Process Regression Modeling

Gaussian Process (GP) regression is well-suited for modeling the complex, non-linear, and non-canonical behaviors typical of bacterial growth data [45]. GP regression accounts for variability arising from intrinsic biological characteristics and experimental conditions, providing a robust framework for predictive modeling and analysis. Specifically, it defines a probability distribution over functions, enabling flexible modeling without assuming a fixed functional form. This non-parametric Bayesian framework describes a prior distribution over functions, which is updated with observed data to yield posterior predictions.

For a given growth curve, the GP prior is constructed using input time points **X** = [**x**_1_, **x**_2_, …, **x**_*n*_] and corresponding optical density (OD) values **Y**. The posterior predictive distribution of OD values at a new input **x** is given by:

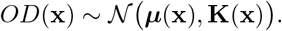

Here, ***µ***(**x**) represents the predicted OD value at a new input **x**, while the covariance **K**(**x**) provides confidence intervals around the prediction. Specifically, **K**(**x**) is defined by the Gaussian kernel:

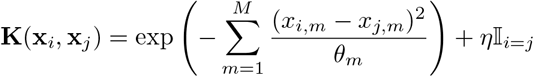

In `DGrowthR`, GP regression is used to model growth curves. To achieve stable and fast GP estimation, we tested several R packages (see Supplementary Table 1) and selected the local approximation GP package `laGP` [46]. `laGP` employs a Gaussian (squared exponential) kernel and is optimized for large-scale spatiotemporal data, enabling `DGrowthR` to be also applicable to long time series data. The Gaussian kernel combined with nugget estimation provided high-quality fits across all datasets analyzed in this study (Supplementary Table 1). Model fitting involves hyperparameter optimization, covariance matrix construction, and OD measurement prediction. Hyperparameters are optimized using gradient-based methods to maximize the marginal log-likelihood. The optimized parameters are then used to construct the covariance matrix **K**, capturing relationships among data points and enabling the model to predict growth at new time points, yielding both mean estimates and uncertainty intervals.

### Growth Parameter Estimation and Statistical Inference

#### Estimation of Growth Parameters

In `DGrowthR`, the modeled response for *OD*(*x*) is used to estimate key growth parameters. From the predicted growth values, the maximum optical density (*OD*_max_), the area under the curve (AUC), and growth loss are directly derived. Since *laGP* does not provide access to a symbolic GP model object, growth rate parameters, including the maximum growth rate (*α*_max_), maximum decay rate (*α*_decay_), and doubling time, are estimated using a high-accuracy finite difference approximation of the first derivative of the Gaussian Process:

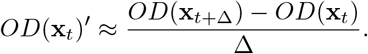

Similarly, phase-transition parameters are approximated using finite difference approximations of the second derivative:

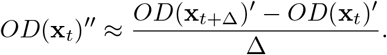

This second derivative approximation also enables the estimation of the lag time *t*_lag_ (the time of greatest increase in growth rate) as well as the time point of entering stationarity *t*_stationary_ (i.e., the time of greatest decrease in growth rate).

#### Differential Growth (DG) Testing

A central capability of `DGrowthR` is testing for significant differences in growth curve dynamics across two distinct conditions. This approach builds on the framework introduced by Tonner et al. [13], which employs a Bayes Factor test statistic to compare the marginal likelihood *p*(**y** | *H*) of the data under two competing hypotheses.

The Bayes Factor is defined as:

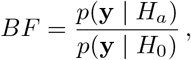

where the null hypothesis (*H*_0_) assumes that all growth curves are identical, modeling OD as a function of time. In this case, each input **x**_*i*_ ∈ **X** is one-dimensional, containing only the time at which the measurement was taken. In contrast, the alternative hypothesis (*H*_*a*_) accounts for differences between conditions by modeling growth curves as a function of both time and an additional covariate. Here, each input **x**_*i*_ becomes two-dimensional, including both the time of measurement and a binary variable indicating the experimental condition [47].

In the context of high-throughput testing, variations in prior probabilities and potential deviations from modeling assumptions motivate the use of an empirical p-value for the observed Bayes Factor, which can be obtained through permutations [48]. In this approach, artificial datasets are generated by randomly reassigning covariate labels to the growth curves while preserving the original distribution of the covariate. Importantly, label shuffling is performed at the level of entire growth curves, ensuring that the temporal structure of the data remains intact. The empirical p-value is then computed as the proportion of permuted Bayes Factors that are greater than or equal to the observed Bayes Factor:

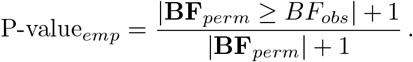

The addition of 1 to both the numerator and denominator prevents the p-value from being exactly zero when none of the permuted Bayes Factors exceed the observed one. This adjustment yields a conservative and unbiased estimate of the p-value, particularly when the number of permutations is limited [49]. In our case, the number of permutations is typically restricted to a maximum of 1000 due to computational constraints, resulting in a minimum achievable p-value of 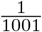 . This resolution is often insufficient to differentiate between highly significant results, as even strong signals will “bottom out” at the minimum possible p-value.

To overcome this limitation, we approximate the distribution of **BF**_perm_ using a Gamma distribution, following the approach proposed by Winkler et al. [20] and implemented in `permApprox` [21]:

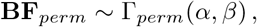

where the shape parameter *α* and the rate parameter *β* are estimated via maximum likelihood estimation. The p-value for the observed Bayes Factor is then derived from the fitted Gamma distribution as follows:

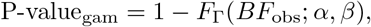

where *F*_Γ_(·; *α, β*) denotes the cumulative distribution function of the Gamma distribution. To save computation time, this approximation is only applied if the empirical p-value P-value_*emp*_ is below 0.2, as our primary interest lies in approximating small p-values as accurately as possible.

### Implementation and Computational Framework

`DGrowthR` is implemented in the R programming language. For a high-level summary of the package, we provide a workflow diagram in Supplementary Fig. 1. The package as well as all code for reproducing the results in this paper are available at https://github.com/bio-datascience/DGrowthR. We next highlight details about specific functionality of the package.

#### Objection-oriented Design and Storage of Results

`DGrowthR` is programmed in an object-oriented fashion and provides the an S4 `DGrowthR` object as fundamental data structure. In addition to performing analyses, `DGrowthR` transparently stores all analysis results, including predicted values, permuted Bayes Factors, and OD differences, within this `DGrowthR` object. These values are accessible for further analysis or integration into external workflows, allowing flexibility for custom downstream analyses.

#### Parallelization and Hyperparameter Summary Statistics

`DGrowthR` leverages parallel computing to enhance efficiency in processing, result compilation, and parameter extraction. With multicore processors and distributed computing environments, `DGrowthR` can concurrently fit and predict models across multiple compounds, significantly reducing computation time. Analytical results are systematically organized and saved as CSV files. Additionally, `DGrowthR` extracts critical hyperparameters from the GP regression fits, such as length scale and nugget parameters, which are essential for assessing model performance and understanding biological implications.

#### Computation and Runtime Analysis

Most computation was conducted on the Helmholtz Munich server using the Linux Ubuntu 20.04 operating system. This server is equipped with 128 CPU cores, 256 threads, and 2 TB of RAM, providing substantial computational resources for handling the present large-scale data analysis tasks. Every `DGrowthR` analysis was computed in parallelized mode with varying number of available cores. To give a realistic assessment of the computational scaling of `DGrowthR` for DG analysis, we also provide an illustrative runtime analysis using the Brenzinger et al. dataset [22]. The results are available in the Supplementary Material E. Briefly, we measured local run times on a MacBook Air with Apple M3 chip (4 performance cores, 4 efficiency cores, 24 GB RAM) using a single core for DG analysis as a function of the number of time points *T* of a growth curve and the number of permutations *n*_perm_ used for inference. The runtime analysis reflects the cubic scaling of GP model fitting with *T* and the linear scaling of the analysis in *n*_perm_ (see Supplementary Fig. 7. At full temporal resolution (*T* = 20 time points in Brenzinger et al. dataset), assessing the statistical significance of a single compound using the default *n*_perm_ = 500 takes ∼30 seconds. Since this computation can be embarrassingly parallelized in `DGrowthR`, the complete analysis of a large-scale dataset can be performed within minutes to hours.

## Data Availability

A complete workflow of `DGrowthR`, including all steps from data pre-processing to analysis and visualization, is available at https://github.com/bio-datascience/DGrowthR. This repository provides a detailed tutorial, a comprehensive vignette, and a sample dataset specifically created for `DGrowthR`. All data used in this study, including the results presented here, are openly accessible, ensuring reproducibility and ease of use. A visualization of the entire workflow is also included to facilitate understanding and implementation of the methods. Publicly available growth data from [23] and [22] were obtained from the respective publications.

## Code Availability

The open-source code for `DGrowthR` is accessible at https://github.com/bio-datascience/DGrowthR. This repository contains the full source code, installation instructions, example scripts, and a visualization of the workflow to assist users in getting started with `DGrowthR`. The code is provided under the MIT License, allowing for broad use and adaptation by the research community.

## Author Contributions

M.F., R.O.A. and C.L.M. conceived the overall objectives and design of the project. M.F., R.O.A., C.L.M., and S.P. contributed to the development of the framework. M.F. and R.O.A. analyzed data from experimental validations and implemented all computational methods. A.Z. and M.K.A. performed the experiments. M.F. and R.O.A. drafted the original manuscript with revisions provided by CLM. All authors revised and approved the final version of the article.

