## Supplementary Material for "Statistical end-to-end analysis of large-scale microbial growth data with DGrowthR"

### A DGrowthR Computational Workflow

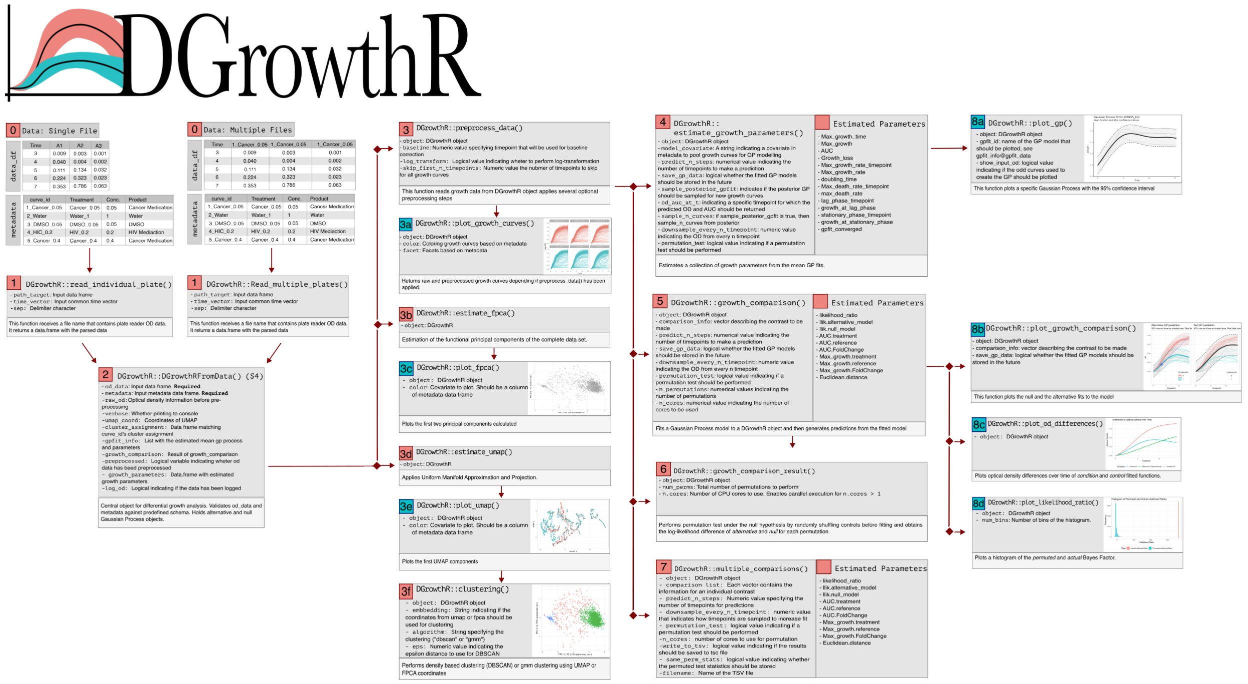


Supplementary Figure 1: Comprehensive workflow of `DGrowthR` Software for Bacterial Growth Analysis. This workflow diagram illustrates the sequential process of utilizing `DGrowthR` for detailed bacterial growth modeling and analysis. The process begins with data import (Step 1), where `DGrowthR` can read either single files or multiple files at once. Next, a `DGrowthR` object is created in Step 2 to handle data and manage subsequent computations. Data can be preprocessed in Step 3, before exploring it further in Step 3a. Functional Principal component analysis can be performed (Step 3b) and visualized (Step 3c),while UMAP coordinates can be calculated (Step 3d) and plotted (Step 3e) to reveal underlying patterns in the data. For grouping similar growth curves, `DGrowthR` offers density-based clustering or Gaussian Mixture modeling (Step 3f). Growth parameters can be automatically estimated in Step 4, while Gaussian Process regression can be applied in Step 5. For statistical analysis, the workflow includes hypothesis testing based on permutations using Step 6 and automated assessment of multiple comparisons (Step 7). Finally, various visualization options (Step 8a-8d), allow for presentation and visual assessment of findings.

#### B Growth Parameter Estimation Strategy

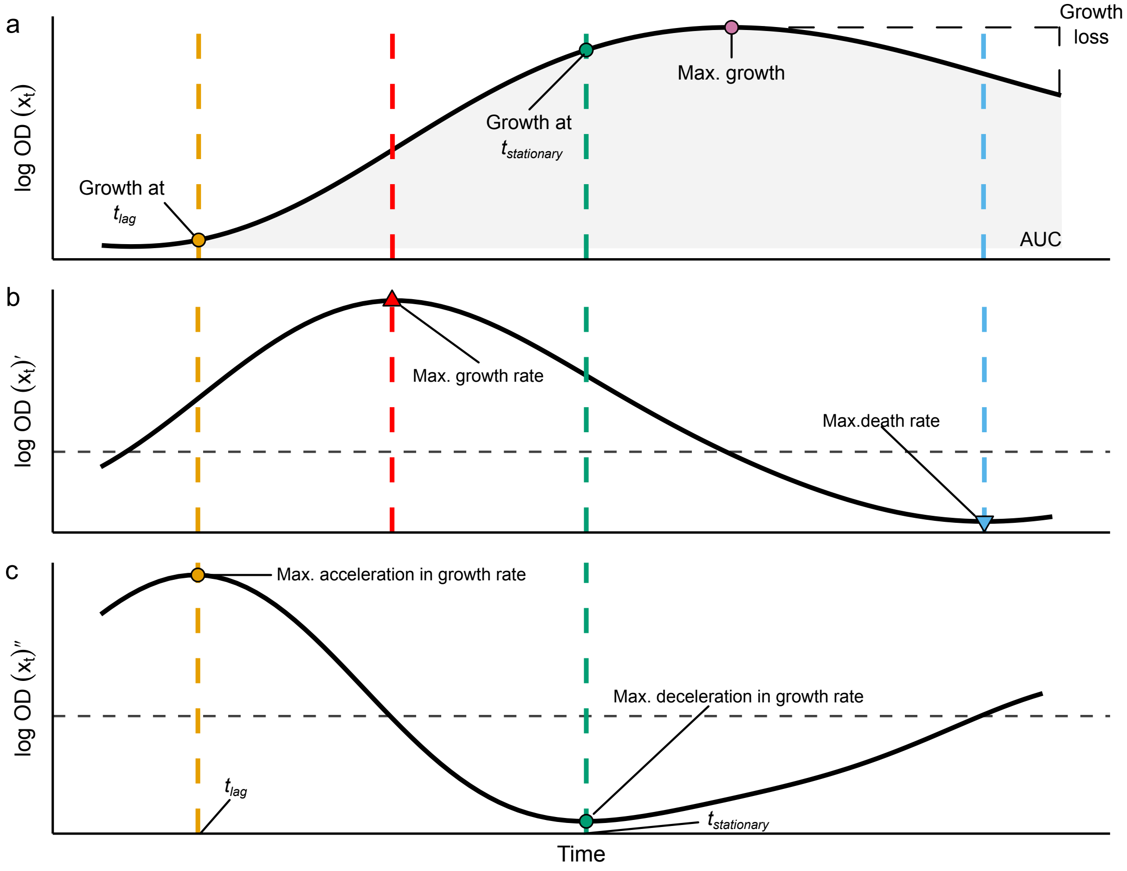


Supplementary Figure 2: Depiction of growth parameter estimation. **a**. GP predicted growth plotted against time. The maximum amount of growth and growth loss are directly observable parameters. Overall growth is recorded as the AUC of the complete predicted growth curve. The amount of growth at the end of the lag phase and the end of the exponential phase is recorded. **b**. The approximated first derivative of the predicted growth. The maximum growth and maximum absolute decay rates are estimated as the maximum and minimum of the first derivative, respectively. The horizontal line indicates 0. **c**. The approximated second derivative of the predicted growth. The time at which the maximum ($t_{\text{lag}}$) and the minimum ($t_{\text{stationary}}$) of the second derivative occur are recorded. This corresponds to the transition between the lag and exponential phases, and the exponential and stationary phases, respectively.

### C Comparison between DGrowthR and Growthcurver for growth curve fitting

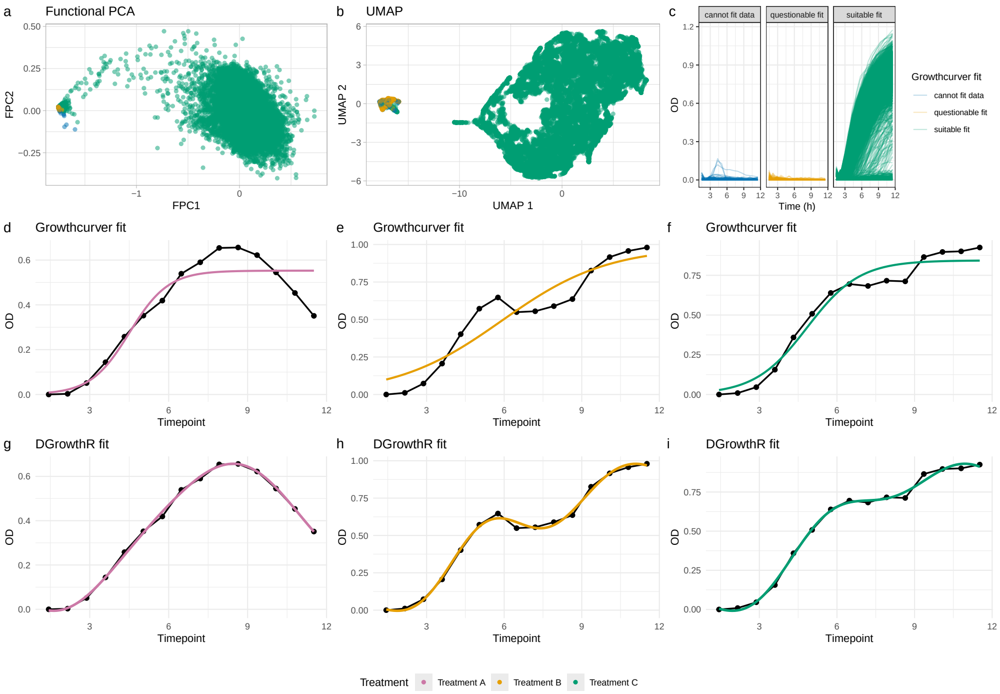


Supplementary Figure 3: Comparison of two frameworks available in R for analyzing bacterial growth curves: `Growthcurver` and `DGrowthR`. **a**. Panel a presents the Functional Principal Component Analysis for *S. enterica*, with points colored according to `Growthcurver`'s fit quality (cannot fit data, questionable fit, suitable fit). **b**.Panel b shows the UMAP projection using the same color scheme, revealing clear separation between suitable fits and the other two categories. **c**.Panel c displays growth curves grouped by fit category, demonstrating that non-growing curves cannot be properly fitted with `Growthcurver` due to its underlying parametric assumptions. **d-f**.Panel d-f present individual growth curves initially classified as "suitable fits" by `Growthcurver`, revealing its limitations. Panel d shows inaccurate estimation of the decay phase, while panels e and f demonstrate poor capture of diauxic growth patters. These limitations stem form the underlying assumption that growth curves follow a sigmoidal pattern. **g-i**.In contrast, Panels g-i illustrate how `DGrowthR` models the complex growth dynamics more accurately. Panel g demonstrates how `DGrowthR` captures the decay phase, while Panels h and i show the ability to model the diauxic growth patterns. The `DGrowthR` framework's flexibility allows it to adapt to diverse bacterial growth patterns without being constrained by parametric assumptions.

### D Dimensionality Reduction and Clustering of Growth Profiles

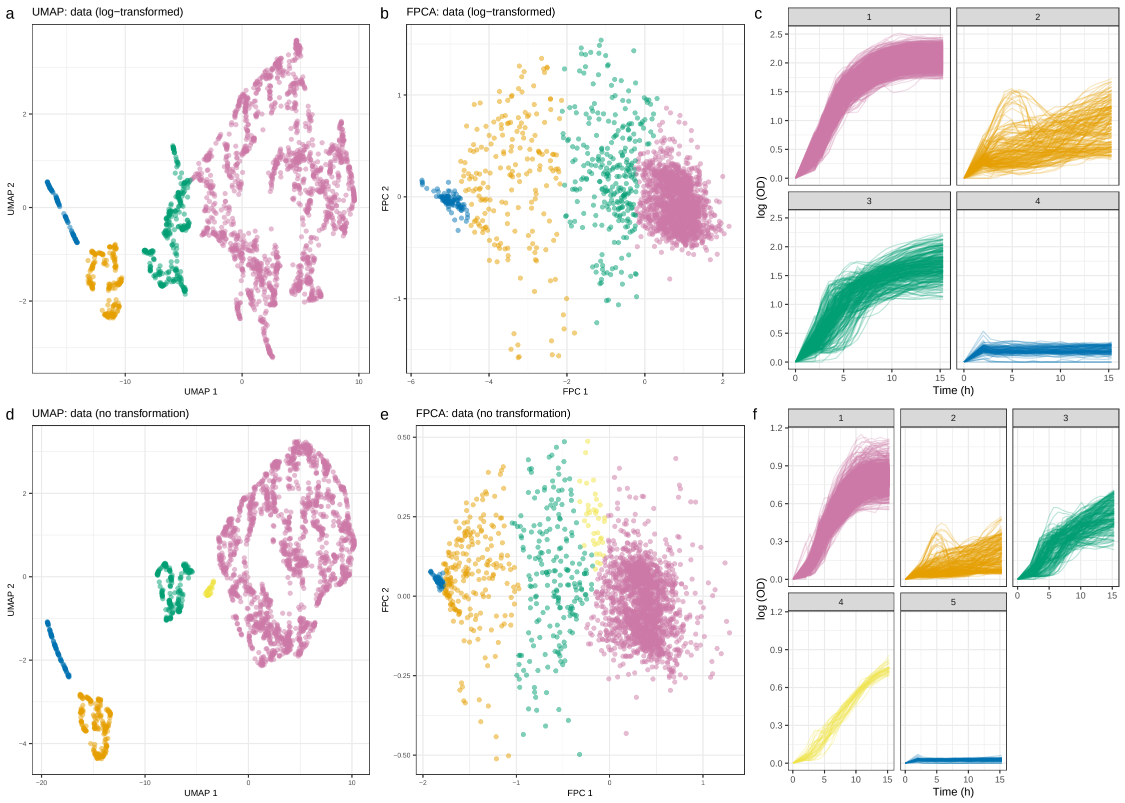


Supplementary Figure 4: Low-dimensional embeddings of the Brenzinger et al. dataset reveal distinct bacterial growth patterns. **a.** Panel a displays the UMAP projection of log-transformed data colored by density-based clustering, highlighting four distinct growth patterns. **b**. Panel b shows the Functional Principal Component Analysis of the same log-transformed data, showing consistent clustering patterns. **c**.Panel c illustrates the corresponding growth curves for each identified cluster. Cluster 1 (pink) shows typical sigmoidal growth reaching high optical density, cluster 4 (blue) shows non-growing curves, while Clusters 2 (orange) and 3 (green) demonstrate intermediate growth with notable deviations from typical sigmoidal patterns. **d**. Panel d shows the UMAP projection of linear (untransformed) data with density-based clustering, which independently identifies five clusters. The overall structure is consistent with the log-transformed analysis, but the finer resolution reveals an additional cluster (Cluster 4, yellow) positioned between Cluster 3 (green) and Cluster 1 (pink). This additional cluster captures curves exhibiting near-linear growth behavior. Both dimensionality reduction techniques demonstrate a consistent gradient from low to high growth moving from left to right across the projection space, observed in both logarithmic and linear transformations of the data.

### E Evaluation of Growth Model Accuracy

To evaluate the necessity of non-parametric modeling for these data, we compared Gaussian Process regression against two widely used parametric growth models [1]. These are:

1. Logistic growth model. The OD at time $t$ ($\hat{y}_t$) is modeled as:

$$\hat{y}_t = \frac{A}{1 + \exp\left(\frac{4\mu_{\max}(\lambda - t)}{A} + 2\right)} \ , \tag{1}$$

   where $A$ is the carrying capacity, $\mu_{\max}$ is the maximum specific growth rate, and $\lambda$ is the duration of the lag phase.

2. Gompertz growth model:

$$\hat{y}_t = A \cdot \exp\left(-\exp\left(\frac{\mu_{\max} \cdot e}{A}(\lambda - t) + 1\right)\right) \ , \tag{2}$$

   where $A$, $\mu_{\max}$, and $\lambda$ have the same meaning as described above.

The parametric models were fit using the `nls` and `nlsLM` [2]functions in R. The carrying capacity ($A$) was initialized as the maximum OD measurement for each growth curve or treatment. The maximum growth rate ($\mu_{\max}$) was initialized as the maximum of the empirical first derivative of each growth curve. The lag phase parameter ($\lambda$) was initialized as the first time point.

In Supplementary Figure 5a, we evaluated the ability of each growth model to model each growth curve. For each curve, model fit quality is quantified using the mean squared error:

$$\frac{1}{n_{\text{tp}}} \sum_{i=1}^{n_{\text{tp}}} (\hat{y}_i - y_i)^2, \tag{3}$$

where $n_{\text{tp}}$ is the total number of measurements made for a given growth curve (here $n_{\text{tp}} = 20$), $y_i$ is the actual OD measurement at index $i$, and $\hat{y}_i$ is the value predicted by a given growth model. For each growth curve, the model with the lowest MSE value is reported as the best-fitting model. Gaussian Process regression outperformed the vast majority of the growth curves, with 96% of all growth curves being best modeled by this approach.

Since DGrowthR jointly models replicate curves for a given treatment, we additionally evaluated model performance at the treatment level. Supplementary Figure 5b shows all replicate growth curves included in the fit of each growth model. For each treatment, we estimated an average MSE by comparing the jointly modeled growth curve to each replicate growth curve:

$$\frac{1}{n_{\text{rep}}} \sum_{j=1}^{n_{\text{rep}}} \frac{1}{n_{\text{tp}}} \sum_{i=1}^{n_{\text{tp}}} (\hat{y}_i - y_{ij})^2, \tag{4}$$

where $n_{\text{rep}}$ is the number of replicate growth curves gathered for a given treatment (typically $n_{\text{rep}} = 3$), $y_{ij}$ is the actual OD measurement at index $i$ for growth curve $j$, and $\hat{y}_i$ is the value predicted by a given growth model. For each treatment, the model with the lowest average MSE is reported as the best performer. Gaussian Process regression is again the best performer across the majority of treatments, with 72% of treatments being best modeled by this approach.

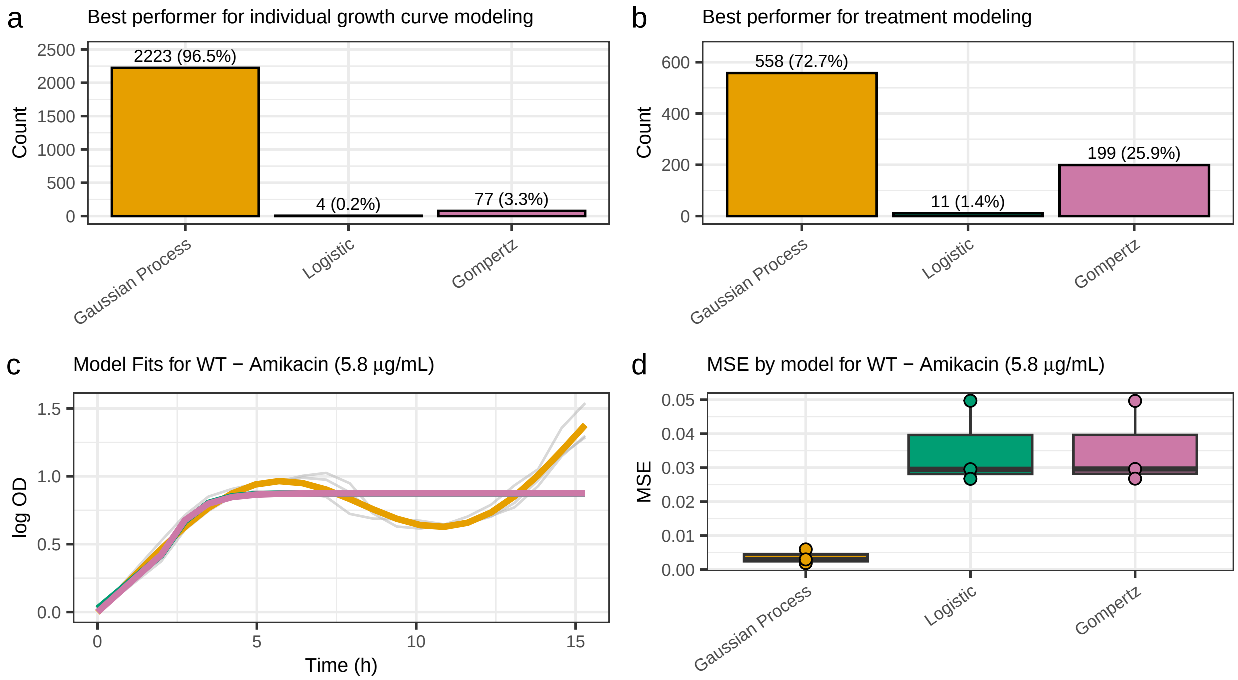


Supplementary Figure 5: Growth model comparison for the Brenzinger et al. (2024) dataset. **a.** Comparison of the number of individual growth curves for which each growth model had the lowest MSE value. **b.** Comparison of the number of joint treatments for which each growth model had the lowest MSE value. **c.** Comparison of model fit for the joint modeling of *V. cholerae* wild-type growth in response to Amikacin (5.8 $\mu$g/mL). Growth curves are shown in grey, model fits follow the color coding of all other panels. **d.** Comparison of MSE values for the individual growth curves gathered for *V. cholerae* wild-type growth in response to Amikacin.

### F Additional illustrations of discovered pairwise interactions

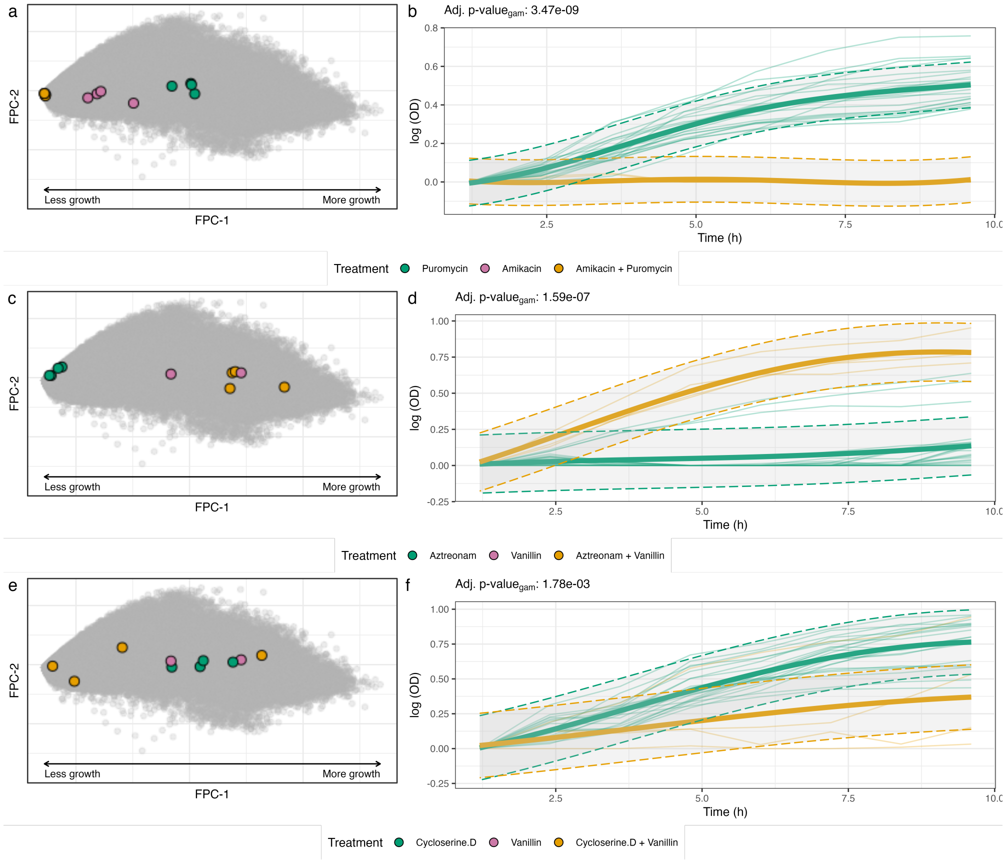


Supplementary Figure 6: Selected pairwise drug interactions highlighting synergistic and antagonistic effects on the growth of *E. coli* BW, identified with `DGrowthR`. For all panels, adjusted gamma-approximated p-values were computed using 500 permutations. **a**. Position of replicate growth curves in the FPCA embedding, gathered from treatment with Puromycin (25 $\mu$g/mL) and Amikacin (0.75 $\mu$g/mL) individually and in combination. **b**. Alternative model of *E. coli* BW growth treated with Puromycin alone and in combination with Amikacin. Shaded area indicates 95% confidence interval. The combination exhibits a strong synergistic effect ($\log_2$FC $= -9.62$, adjusted p-value$_{\text{gam}}$ $= 3.47 \times 10^{-9}$), representing the lowest AUC fold change among all tested combinations. **c**. Position of replicate growth curves in the FPCA embedding, gathered from treatment with Aztreonam (0.06 $\mu$g/mL) and Vanillin (200 $\mu$g/mL) individually and in combination. **d**. Alternative model of *E. coli* BW growth treated with Aztreonam alone and in combination with Vanillin. Shaded area indicates 95% confidence interval. The combination exhibits the highest positive AUC fold change among all tested combinations ($\log_2$FC $= 3.03$, adjusted p-value$_{\text{gam}}$ $= 1.59 \times 10^{-7}$), indicating an antagonistic effect consistent with the general antagonism of Vanillin against antibiotics reported by Brochado et al. [3]. **e**. Position of replicate growth curves in the FPCA embedding, gathered from treatment with Cycloserine D (8 $\mu$g/mL) and Vanillin (200 $\mu$g/mL) individually and in combination. **f**. Alternative model of *E. coli* BW growth treated with Cycloserine D alone and in combination with Vanillin. Shaded area indicates 95% confidence interval. A modest but significant reduction in growth is observed ($\log_2$FC $= -1.07$, adjusted p-value$_{\text{gam}}$ $= 1.78 \times 10^{-3}$).

### G Gaussian Process Library Selection

The differential growth testing framework in `DGrowthR` places specific requirements on the underlying Gaussian Process library. First, a nugget parameter is needed to ensure numerical stability of the covariance matrix, as growth data frequently yield matrices that are not positive definite. Second, access to the marginal log-likelihood is essential for computing the Bayes Factor test statistic. Third, the alternative model in the hypothesis testing framework models growth as a function of both time and experimental condition, requiring support for multiple covariates with anisotropic length scales. We evaluated six Gaussian Process libraries available in R and Python against these requirements (Supplementary Table 1).

| **Feature** | **kernlab** | **DiceKriging** | **Gpfit** | **Kergp** | **Gpy** | **laGP** |
|---|---|---|---|---|---|---|
| Language | R | R | R | R | Python | R |
| Nugget | ✓ | × | ✓ | ✓ | ✓ | ✓ |
| Optimization | × | × | ✓ | ✓ | × | ✓ |
| Different Kernels | ✓ | ✓ | ✓ | ✓ | ✓ | × |
| Log-Likelihood | × | ✓ | Limited | ✓ | ✓ | ✓ |
| Multiple Covariates | ✓ | ✓ | ✓ | ✓ | ✓ | ✓ |
| Anisotropic Models | × | × | × | ✓ | ✓ | ✓ |

Supplementary Table 1: Comparison of Gaussian Process libraries evaluated for use in `DGrowthR`. Features critical for the differential growth testing framework include nugget support, hyperparameter optimization, log-likelihood access, multiple covariates, and anisotropic modeling. Among the R packages, only `laGP` [4] satisfies all requirements simultaneously. While `kergp` [5] also supports these features, it exhibited numerical instability on growth curve data. `GPy` [6] is included as a reference from the Python ecosystem. The primary limitation of `laGP` is the restriction to a Gaussian (squared exponential) kernel. However, this proved sufficient across all datasets analyzed in this study.

### H Computational Runtime Analysis

Supplementary Figure 7 depicts the per-contrast computational performance of `DGrowthR` using runtime benchmarking. All experiments were performed locally on a MacBook Air with Apple M3 chip (4 performance cores, 4 efficiency cores, 24 GB RAM) using a single core. The Brenzinger et al. [7] dataset ($n = 378$ contrasts) was used for benchmarking. We evaluated the performance across varying permutation counts ($n_{\text{perm}} \in \{10, 50, 100, 500, 1000\}$) and downsampling factors (1, 2, and 3), corresponding to $T = 20$, 10, and 7 effective timepoints, respectively. A downsampling factor of 1 represents the full analysis without data reduction.

To characterize the scaling behavior, we first confirmed that runtime scales linearly with $n_{\text{perm}}$ at each temporal resolution by fitting separate per-timepoint regressions ($R^2 > 0.986$ in all cases). We then fitted a reduced linear regression model incorporating the cubic dependence on $T$:

$$t = \beta_0 + \beta_1 \, n_{\text{perm}} + \beta_2 \, n_{\text{perm}} \cdot T^3 \tag{5}$$

where $t$ is the per-contrast runtime in seconds, $n_{\text{perm}}$ is the number of permutations, and $T$ is the effective number of timepoints. The $T^3$ term reflects the $\mathcal{O}(T^3)$ complexity of Gaussian Process covariance matrix inversion, which dominates the cost of each permutation. The main effect of $T^3$ (without interaction) was not significant ($p = 0.164$, $F$-test) and was therefore excluded from the final model. This is expected, since in the absence of permutations ($n_{\text{perm}} = 0$), no GP model is fitted, and hence the number of timepoints does not contribute to the runtime. The cubic scaling of $T$ depends only on the permutation which is precisely what the interaction term $\beta_2 \, n_{\text{perm}} \cdot T^3$ captures.

The final model achieved $R^2 = 0.979$ with a residual standard error of 2.44 s. All coefficients were highly significant ($p < 2 \cdot 10^{-16}$):

$$\hat{\beta}_0 = 1.031 \quad \text{(baseline overhead per contrast)} \tag{6}$$

$$\hat{\beta}_1 = 3.963 \times 10^{-3} \quad \text{(per-permutation cost independent of } T\text{)} \tag{7}$$

$$\hat{\beta}_2 = 7.504 \times 10^{-6} \quad \text{(cubic scaling coefficient)} \tag{8}$$

The predicted mean runtimes closely matched the observed values across all 15 experimental conditions (Table 2). The effective per-permutation cost grows from approximately 6.5 ms at $T = 7$ to 64.1 ms at $T = 20$, consistent with cubic scaling. This explains the practical effectiveness of downsampling. Even

Supplementary Table 2: Observed and predicted mean per-contrast runtimes (seconds). All residuals are below 0.6 seconds in absolute terms.

| $n_{\text{perm}}$ | $T$ | Observed (s) | Predicted (s) | Residual (s) |
|---|---|---|---|---|
| 10 | 7 | 1.0 | 1.1 | −0.1 |
| 10 | 10 | 1.1 | 1.1 | 0.0 |
| 10 | 20 | 1.6 | 1.7 | 0.0 |
| 50 | 7 | 1.2 | 1.4 | −0.1 |
| 50 | 10 | 1.6 | 1.6 | 0.0 |
| 50 | 20 | 4.2 | 4.2 | −0.1 |
| 100 | 7 | 1.8 | 1.7 | 0.2 |
| 100 | 10 | 2.1 | 2.2 | 0.0 |
| 100 | 20 | 7.4 | 7.4 | −0.1 |
| 500 | 7 | 4.9 | 4.3 | 0.6 |
| 500 | 10 | 6.8 | 6.8 | 0.0 |
| 500 | 20 | 32.9 | 33.0 | −0.1 |
| 1000 | 7 | 7.3 | 7.6 | −0.3 |
| 1000 | 10 | 12.4 | 12.5 | −0.1 |
| 1000 | 20 | 65.1 | 65.0 | 0.1 |

the slowest downsampled configuration ($T = 7$, $n_{\text{perm}} = 1000$; 7.3 s) completed faster than the fastest full-resolution configuration at equivalent permutation depth ($T = 20$, $n_{\text{perm}} = 1000$; 65.1 s). Based on these estimates, the Brochado et al. [3] analysis (5,795 comparisons, 500 permutations, $T = 20$) would require approximately 53 hours on a single core, reducing to roughly 7 hours when parallelized across 8 cores.

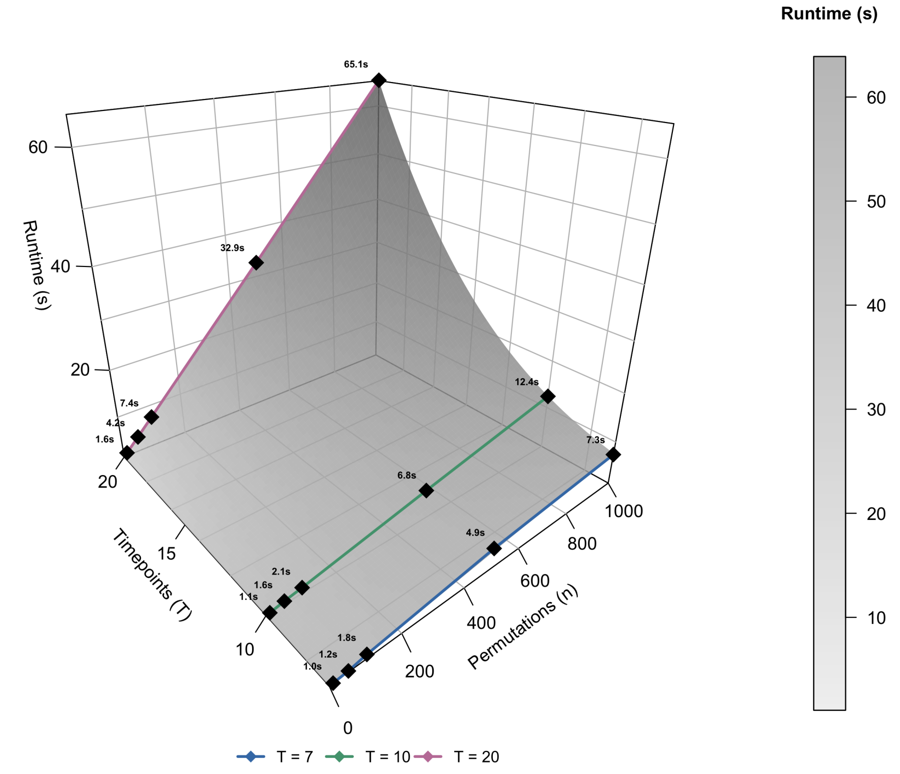


Supplementary Figure 7: Computational scalability of `DGrowthR`. Three-dimensional surface plot showing the predicted per-contrast runtime (seconds) as a function of the number of permutations ($n_{\text{perm}}$) and effective timepoints ($T$), based on the fitted model $t = \hat{\beta}_0 + \hat{\beta}_1\, n_{\text{perm}} + \hat{\beta}_2\, n_{\text{perm}} \cdot T^3$ ($R^2 = 0.979$). The grey surface represents the model prediction. Colored lines connect the observed mean runtimes across permutation counts for each temporal resolution: $T = 7$ (blue), $T = 10$ (green), and $T = 20$ (pink). Black diamonds mark the observed means, annotated with their values in seconds. The steep rise of the surface along the $T$-axis at high $n_{\text{perm}}$ reflects the $\mathcal{O}(T^3)$ complexity of the underlying Gaussian Process matrix operations. All experiments were performed on a single core of a MacBook Air (Apple M3, 24 GB RAM) using the Brenzinger et al. [7] dataset ($n = 378$ contrasts).

[5] Yves Deville, David Ginsbourger, Olivier Roustant Contributors, Nicolas Durrande, and Maintainer Olivier Roustant. *kergp: Gaussian Process Laboratory*, 2015.

[6] GPy. GPy: A gaussian process framework in python. `http://github.com/SheffieldML/GPy`, since 2012.

[7] Susanne Brenzinger, Martina Airoldi, Adewale Joseph Ogunleye, Karl Jugovic, Martin Krähenbühl Amstalden, and Ana Rita Brochado. The vibrio cholerae cbass phage defence system modulates resistance and killing by antifolate antibiotics. *Nature Microbiology*, 9(1):251–262, January 2024. ISSN 2058-5276. doi: 10.1038/s41564-023-01556-y.
